# Nuclear GAPDH cascade mediates stress-dependent pathological cardiac growth

**DOI:** 10.1101/844902

**Authors:** Shun Nakamura, Manling Zhang, Genri Numata, Taro Kariya, Hideyuki Sasaki, Masayuki Sasaki, Adriana Ramos, Eisuke Amiya, Masaru Hatano, Masahiko Ando, Yoichi Yasunaga, Masaki Iwakiri, Norimichi Koitabashi, Guangshuo Zhu, Tsuyoshi Tsujimura, Dong I. Lee, Carlos Tristan, Neelam Shahani, Yukihiro Tsuchiya, Hanna Jaaro-Peled, Tetsuo Sasano, Barbara Slusher, David A. Kass, Tetsuo Ushiku, Minoru Ono, Toshiaki Saitoh, Koko Ishizuka, Akira Sawa, Eiki Takimoto

**Affiliations:** Department of Cardiovascular Medicine, The University of Tokyo School of Medicine, Tokyo, Japan; Department of Cardiovascular Medicine, Institute of Science Tokyo, Tokyo, Japan; Division of Cardiology, Johns Hopkins University School of Medicine, Baltimore, MD, USA; Isotope Science Center, The University of Tokyo, Tokyo, Japan; Department of Anesthesiology, The University of Tokyo School of Medicine, Tokyo, Japan; Department of Psychiatry, Johns Hopkins University School of Medicine, Baltimore, MD, USA; Advanced Medical Center for Heart Failure, The University of Tokyo Hospital, Tokyo, Japan; Department of Cardiac Surgery, The University of Tokyo, Tokyo, Japan; Department of Pathology, The University of Tokyo, Tokyo, Japan; Department of Pharmacology, Showa Pharmaceutical University, Tokyo, Japan; Department of Neurology, Johns Hopkins University School of Medicine, Baltimore, MD, USA; Department of Neuroscience, Johns Hopkins University School of Medicine, Baltimore, MD, USA; Department of Pharmaceutical Sciences, Nihon Pharmaceutical University, Saitama, Japan; Department of Biomedical Engineering, Johns Hopkins University School of Medicine, Baltimore, MD, USA; Department of Pharmacology, Johns Hopkins University School of Medicine, Baltimore, MD, USA; Department of Genetic Medicine, Johns Hopkins University School of Medicine, Baltimore, MD, USA; Department of Mental Health, Johns Hopkins University Bloomberg School of Public Health, Baltimore, MD, USA

## Abstract

Here we report that stress-induced nuclear translocation of GAPDH mediates heart hypertrophy via Brahma-Related-Gene-1 (BRG1)-associated chromatin remodelling. In response to pressure overload elicited by transverse aortic constriction, we observed nuclear translocation of GAPDH in the mouse heart. We also demonstrated a robust nuclear localization of GAPDH in cardiomyocytes from patients with dilated hypertrophic cardiomyopathy, whereas negligible GAPDH in the nucleus in control subjects. This is the first demonstration of disease-associated nuclear GAPDH directly in living patients. Using immunohistochemical methods and a pharmacological way that selectively blocks GAPDH nuclear translocation (RR compound), we proved the causal involvement of GAPDH cysteine-150 modification in this translocation in Gq-induced cell model for heart hypertrophy. Accordingly, both pharmacological and genetic interventions proved that the same mechanism played a causal role for heart hypertrophy and dysfunction *in vivo*. We discovered that, upon nuclear translocation, GAPDH augmented the protein interaction of BRG1 and histone deacetylase 2 (HDAC2), which further facilitated the *Myh7/Myh6* isoform ratio from the mature to immature status, an essential mechanism for heart hypertrophy. Beyond medical implications, we provide a novel mechanism of stress-induced reversion of a cellular phenotype from adult to fetal state, which is mediated by a “moonlighting” function of GAPDH.

## Main

Glyceraldehyde-3-phosphate dehydrogenase (GAPDH) has multiple roles in diverse cellular processes beyond its role as a glycolytic enzyme^1–15^, currently acknowledged as one of the most representative “moonlighting” proteins^1,16,17^. One of the most well-studied non-glycolytic cellular features associated with GAPDH is its nuclear translocation in response to various stressors [nuclear GAPDH (N-GAPDH) cascade]^13,18–20^. Although originally discovered in neuronal cells, this translocation is known to occur in both neuronal and non-neuronal cells^13,18,19^.

More than one type of posttranslational modification on GAPDH functions as a trigger for this translocation, which results in interactions between GAPDH and nuclear proteins, including histone acetyltransferase p300 and CREB-binding protein (CBP), as well as NAD^+^-dependent deacetylase SIRT1^19,20^. The nuclear partners of GAPDH are likely to be context-dependent; as evidenced by the interaction with SIRT1 in cell autophagic induction^20^, versus the interaction with p300 in cell death^19^. Investigators have hypothesized that altered histone-related nuclear signalling elicited by N-GAPDH affect gene transcription, which may impact the cellular outcome^2^. Nevertheless, the overall landscape of N-GAPDH cascade remains elusive. The direct link between the regulation of GAPDH with its nuclear protein interactors and specific gene transcription for the final phenotype has not been convincingly shown yet. Furthermore, a major knowledge gap is that N-GAPDH has not been directly demonstrated in human pathological conditions, in contrast to healthy conditions, *in vivo*.

Heart hypertrophy is a central medical problem in cardiology^21^. Models of heart hypertrophy have been frequently built by extrinsic stressors, such as transverse aortic constriction (TAC) for mouse models and Gq-agonists for cell models^22–26^. Intracellularly, heart remodelling that underlies pathological hypertrophy is regulated by the protein interaction of histone deacetylase 2 (HDAC2) and Brahma-Related-Gene-1 (BRG1)^27,28^. This protein interaction is responsible for an isoform shift from adult-type myosin heavy chain [*myosin, heavy polypeptide 6, cardiac muscle, alpha* (*Myh6*)] to its fetal form *Myh7*, leading to cardiomyocyte remodelling. The mechanism whereby extrinsic stressors can impact the HDAC-associated nuclear changes remains elusive.

### N-GAPDH in mouse heart hypertrophy

Our group have used a mouse model of pressure-overload heart hypertrophy induced by TAC to study the mechanism of heart hypertrophy^22–24^, in which the involvement of extrinsic stressors and HDAC-associated nuclear changes have been suggested^29^. Given that N-GAPDH cascade may possibly link these two, we tested whether stress-induced GAPDH nuclear translocation (activation of N-GAPDH cascade) occurs in this model. Heart hypertrophy developed 10 days after TAC (**Fig. 1a**). We examined GAPDH by immunostaining in isolated cardiomyocytes from TAC hearts and observed nuclear GAPDH (**Fig. 1b**). This nuclear translocation of GAPDH was further confirmed by protein subcellular fractionation analysis of left ventricular tissues, while the difference in the levels of cytosolic GAPDH was negligible between sham and TAC hearts (**Fig. 1c,d**). This observation is in accordance with the past reports on N-GAPDH cascade in neurons and other cells, in which a small fraction of cytosolic GAPDH is utilized for N-GAPDH cascade under stressors in a gain-of-function manner^13,30^. No sex difference between males and females was observed regarding the nuclear translocation of GAPDH (**Extended Data Fig. 1**). The change in mRNA expression of *BCL2 binding component 3* (*Bbc3*, encoding PUMA) and *BCL2-associated X protein* (*Bax*), which are reportedly changed when nuclear GAPDH translocation mediates cell death^19^, was negligible (**Fig. 1e**), indicating no involvement of apoptosis in this context. Altogether, we show involvement of the N-GAPDH cascade in TAC stress-induced heart hypertrophy, in which no robust cell death is involved.

**Fig. 1:**
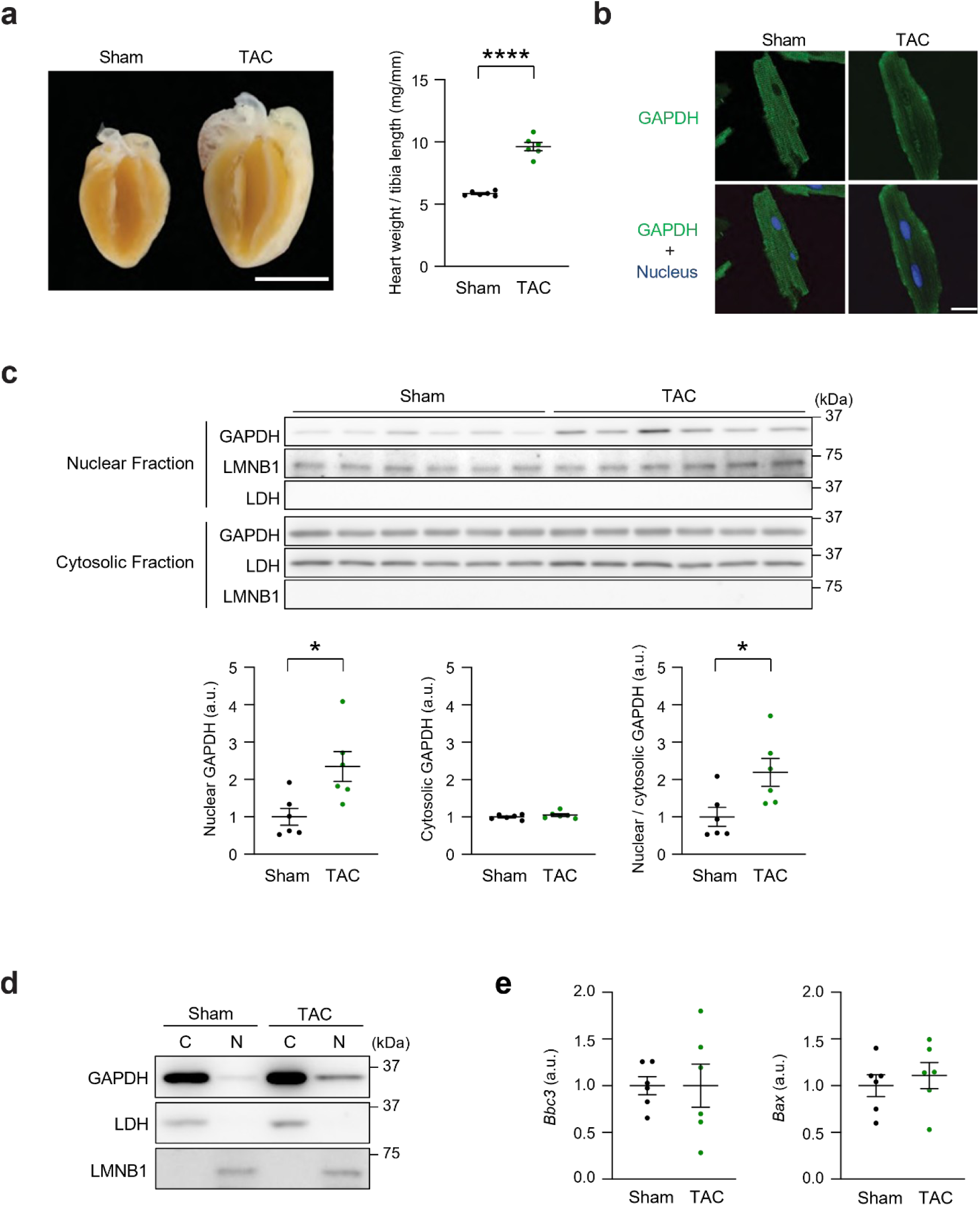
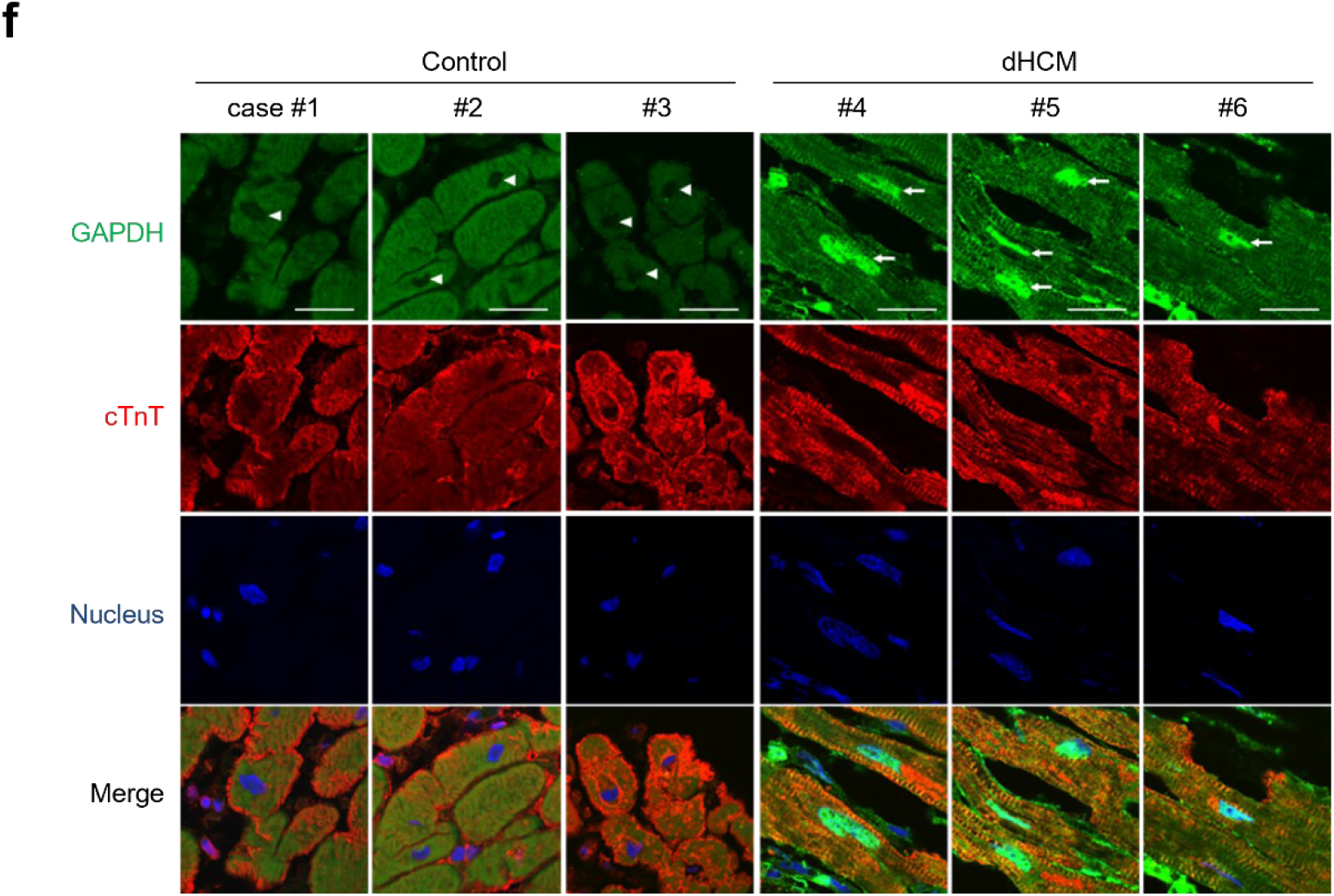
Nuclear translocation of GAPDH is observed in mouse and human heart hypertrophy. **a**, Representative images of sham and TAC hearts (left). Heart weight normalized to tibia length indicates ∼40% increase in TAC hearts compared with sham hearts. Scale bar, 5 mm. Data are expressed as mean ± standard error of the mean (SEM) (*n* = 6 animals per group). An unpaired two-tailed Student’s *t* test was performed. \*\*\*\**p* < 0.0001. **b**, Demonstration of nuclear GAPDH in cardiomyocytes from TAC hearts, but not in those from sham hearts in immunofluorescence staining. Scale bar, 10 µm. **c**, Confirmation of GAPDH nuclear translocation in TAC but not in sham hearts by immunoblots using nuclear and cytosolic fractionated proteins (upper) and quantification results (lower). Lamin B1 (LMNB1) and lactate dehydrogenase (LDH) for nuclear and cytosolic loading controls, respectively. Quantification results are presented as mean ± SEM (*n* = 6 animals per group). Unpaired two-tailed Student’s *t* tests were performed. \**p* < 0.05. **d**, Demonstration of a small but significant amount of GAPDH in the TAC nucleus. C, cytosolic fraction; N, nuclear fraction. The protein amount loaded in the C fraction is one tenth of that of the N fraction, which allows for the quantitative comparison between cytosolic and nuclear GAPDH. **e**, Unaltered mRNA expression levels of *Bbc3* and *Bax* in TAC hearts vs. sham hearts. Data are presented as mean ± SEM (*n* = 6 animals per group). Unpaired two-tailed Student’s *t* tests were performed. **f**, Nuclear localization of GAPDH in cardiomyocytes from patients with dHCM, whereas negligible GAPDH in the nucleus in control subjects, shown by immunofluorescence staining of GAPDH. Arrows, robust accumulation of GAPDH in the nucleus; arrowheads, negligible levels of nuclear GAPDH. Cardiac troponin T (cTnT) marking cytoplasm of cardiomyocytes. Scale bar, 20 µm.

### N-GAPDH in human heart hypertrophy *in vivo*

Stress-induced, nuclear-translocated GAPDH in pathological conditions has been studied, called as N-GAPDH cascade, over the past two decades. Nevertheless, no study has directly demonstrated this nuclear translocation in any human disease condition. To fill this gap, we obtained myocardial tissue from living patients with dilated hypertrophic cardiomyopathy (dHCM) during their surgical implantation of a left ventricular assist device. We chose this condition because it has a clinical trajectory from a hypertrophied phase to a dilated phase, which is equivalent to the pathophysiology of the mouse TAC model.

We examined the biopsied tissue by immunohistochemistry. We observed a robust nuclear localization of GAPDH in cardiomyocytes from human patients with dHCM. Control cardiomyocytes were obtained from heart transplant recipients with no histological signs of rejection. In contrast to dHCM hearts, nuclear GAPDH was negligible in all control hearts (**Fig. 1f, Supplementary Table 1**).

### Intervention of the pathology by the RR compound

One of the main mechanisms that trigger stress-induced nuclear GAPDH translocation is posttranslational modification of cysteine-150 (C150) in GAPDH^13,19^. C150 posttranslational modification of GAPDH enables the protein to interact with Siah1 and translocate to the nucleus^13^. We have developed a compound called RR that selectively blocks GAPDH from interacting with its nuclear chaperon Siah1 via the C150 posttranslational modification-mediated mechanism^30^. Importantly, the RR compound does not affect the glycolytic function of GAPDH^30^, and exhibited no significant interaction with primary targets, including receptors, channels, transporters, or enzymes (**Supplementary Table 2**).

Gq-agonists, such as endothelin-1 (ET-1), angiotensin II (AngII), or phenylephrine (PE), are frequently used as triggers for cell models of heart hypertrophy^22,23,25,26,31^. In cultured cardiomyocytes treated with these Gq-agonists, nuclear GAPDH translocation was consistently observed (**Fig. 2a**). Thus, using these cell models, we tested whether the RR compound blocks the nuclear translocation of GAPDH in cardiomyocytes and observed its robust blockade (**Fig. 2b**). Sulphonated GAPDH is known to selectively accumulate in the nucleus when its nuclear translocation occurs with C150 posttranslational modification^13,19^. We observed sulphonated GAPDH in the nucleus of stressed cardiomyocytes, which was completely inhibited by RR (**Fig. 2c**). RR attenuated ET-1-induced cellular hypertrophic responses, such as an increased cell surface area, protein synthesis assessed by ^3^H-leucine uptake, and *natriuretic peptide type B* (*Nppb*, encoding B-type natriuretic peptide) mRNA expression (**Fig. 2d**). Altogether, these pharmacological data support the role of stress-induced nuclear GAPDH translocation in Gq-agonist-elicited cardiac cellular hypertrophy.

**Fig. 2:**
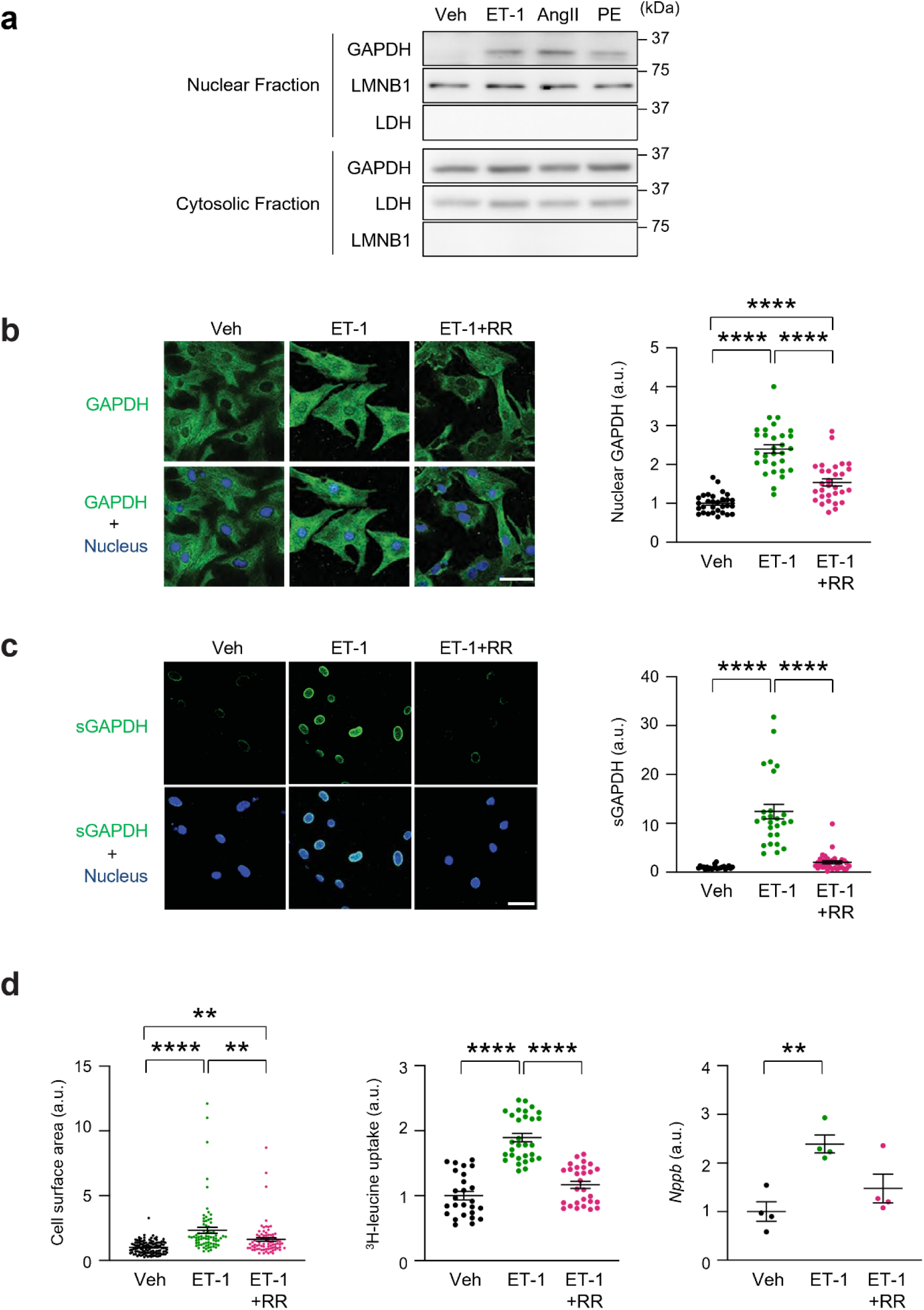
Pharmacological blockade of N-GAPDH cascade with RR attenuates hypertrophy in a heart cellular model. **a**, Nuclear GAPDH in H9c2 cells stimulated with ET-1, AngII, PE, whereas no nuclear GAPDH in cells with vehicle (Veh), shown by immunoblots for subcellular fractionated proteins. LMNB1 and LDH for nuclear and cytosolic loading controls, respectively. **b**, Blockade of N-GAPDH by the RR compound in isolated rat neonatal cardiomyocytes exposed to Veh, ET-1, or ET-1+RR in immunofluorescence imaging (left) and the quantification results using the nuclear-to-cytosolic intensity ratio of GAPDH (right). Scale bar, 50 µm. Data are presented as mean ± SEM (*n* = 30 cells per group). One-way analysis of variance (ANOVA) followed by Bonferroni multiple comparison tests were performed. \*\*\*\**p* < 0.0001. **c**, Sulphonated GAPDH (sGAPDH) in the nucleus, the evidence that this nuclear translocation is mediated by GAPDH C150 modification, in isolated rat neonatal cardiomyocytes exposed to ET-1. Its inhibition by the RR compound is shown in immunofluorescence imaging (left). The quantification results for sGAPDH intensity are also shown (right). Data are presented as mean ± SEM (*n* = 23, 26, and 33 cells for Veh, ET-1, and ET-1+RR, respectively). One-way ANOVA followed by Bonferroni multiple comparison tests were performed. \*\*\*\**p* < 0.0001. **d**, Inhibition of ET-1-induced cellular hypertrophy with the RR compound, assessed by cell surface area (*n* = 114, 74, and 78 cells for Veh, ET-1, and ET-1+RR, respectively), ^3^H-leucine uptake (*n* = 26, 30, and 28 wells for Veh, ET-1, and ET-1+RR, respectively), and *Nppb* mRNA expression (*n* = 4 biologically independent preparations per group). Data are presented as mean ± SEM. One-way ANOVA followed by Bonferroni multiple comparison tests were performed. \*\**p* < 0.01 and \*\*\*\**p* < 0.0001.

Based on these cell data, we employed RR in the TAC model. Daily treatment with RR (0.25 mg/kg/day i.p.) from the beginning of TAC induction reduced nuclear GAPDH in TAC hearts (**Fig. 3a**) and attenuated hypertrophic remodelling of the hearts, as indicated by reduced heart weight, cardiomyocyte size, and fibrosis area (**Fig. 3b,c**). RR treatment mitigated the re-induction of *Myh7* and other gene expression profiles [*Nppb*, *ATPase, Ca^++^ transporting, cardiac muscle, slow twitch 2 (Atp2a2)*, and *phospholamban* (*Pln)*], all of which are associated with heart hypertrophy at the molecular levels (**Fig. 3d**). RR treatment also improved cardiac function and chamber size in TAC hearts, as reflected by increased fractional shortening (FS) and decreased left ventricular end-diastolic dimension (LV-EDD) (**Fig. 3e**). Additionally, an invasive hemodynamic study using pressure-volume (PV) loop analysis revealed that RR treatment enhanced cardiac systolic function (dP/dtmax: maximum first derivatives of left ventricular pressure [LVP]) and improved diastolic function (Tau), under elevated pressure overload, as indicated by arterial elastance (Ea) and peak LVP (**Fig. 3f**). These findings indicate that RR effectively inhibits the N-GAPDH cascade *in vivo*, preventing pathological molecular changes such as *Myh7* induction and overall heart hypertrophy in the TAC model.

**Fig. 3:**
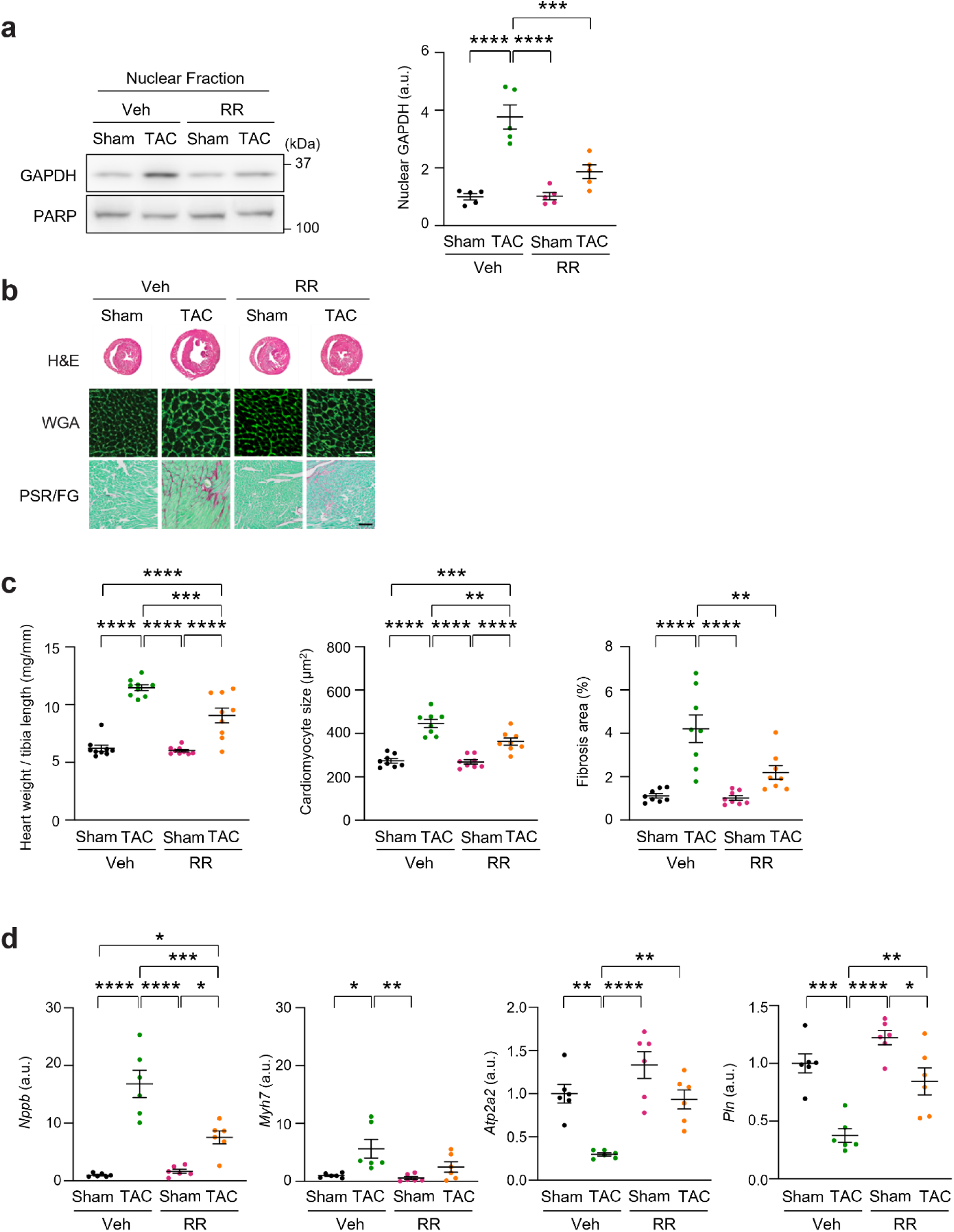
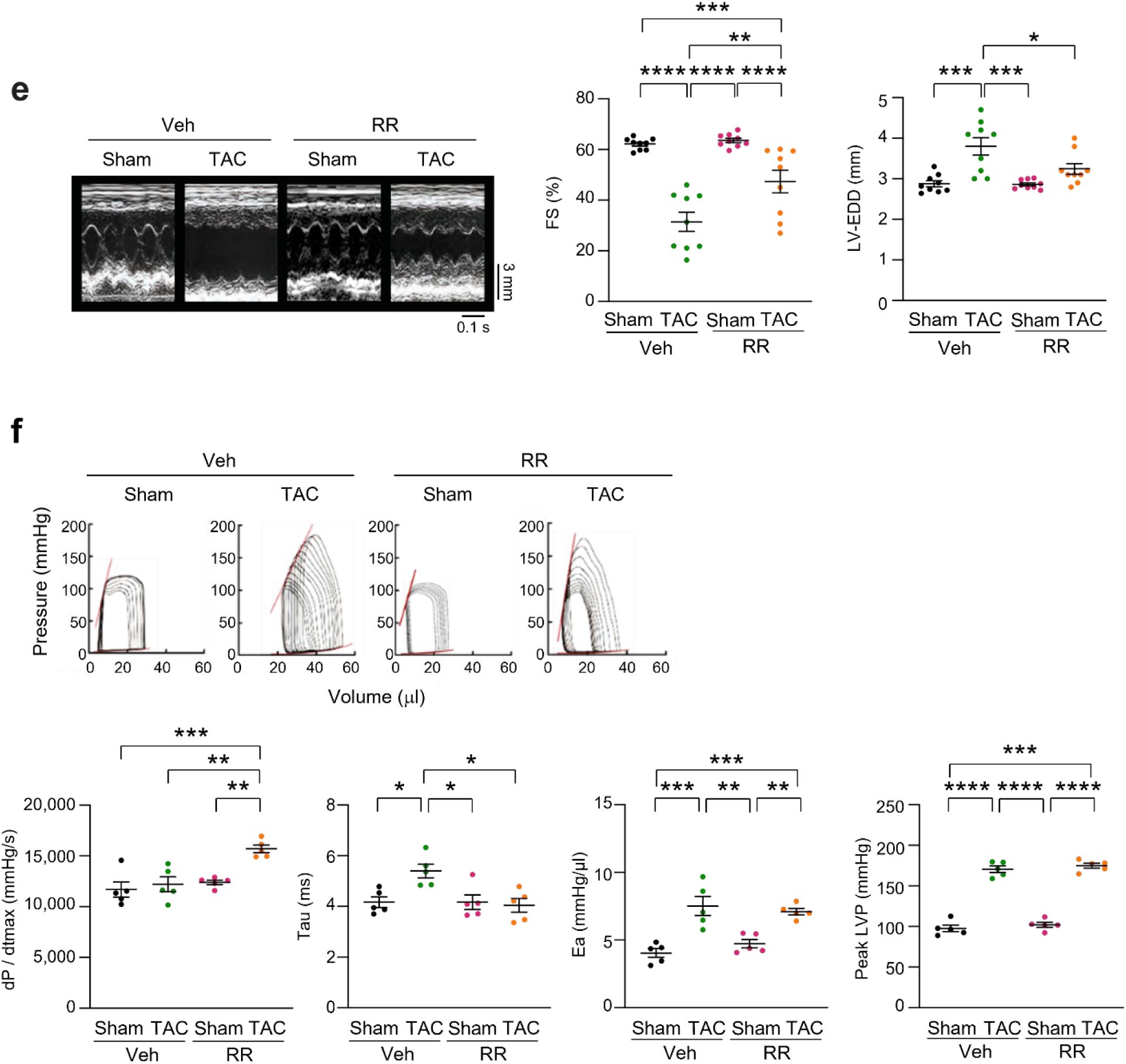
Pharmacological inhibition of N-GAPDH cascade with RR ameliorates heart hypertrophy in mouse TAC model. **a**, Inhibition of nuclear translocation of GAPDH in TAC hearts by the RR compound. Immunoblots of nuclear fractionated proteins (left) and the quantification results (right) are shown. TAC- or sham-operated mice were treated with Veh or RR daily after surgery. Poly (ADP-ribose) polymerase (PARP) for a loading control. Data are presented as mean ± SEM (*n* = 5 animals per group). Two-way ANOVA followed by Bonferroni multiple comparison tests were performed. \*\*\**p* < 0.001 and \*\*\*\**p* < 0.0001. **b**, Inhibition of TAC-induced heart hypertrophy with fibrosis by the RR compound. Representative images of haematoxylin and eosin (H&E), wheat germ agglutinin (WGA), and Picro Sirius Red/Fast Green (PSR/FG) staining of heart cross-sections. Scale bars, 3 mm (H&E); 50 µm (WGA); and 100 µm (PSR/FG). **c**, RR inhibition of TAC-induced heart hypertrophy and fibrosis, assessed by heart weight normalized to tibia length, cardiomyocyte size measured from WGA-stained sections, and fibrosis area assessed using PSR/FG staining. Data are presented as mean ± SEM (*n* = 9, 8, and 8 animals per group for heart weight normalized to tibia length, cardiomyocyte size, and fibrosis area, respectively). Two-way ANOVA followed by Bonferroni multiple comparison tests were performed. \*\**p* < 0.01, \*\*\**p* < 0.001, and \*\*\*\**p* < 0.0001. **d**, Amelioration of myocardial mRNA expression profiles for *Nppb*, *Myh7*, *Atp2a2,* and *Pln* by the RR compound. Data are presented as mean ± SEM (*n* = 6 animals per group). \**p* < 0.05, \*\**p* < 0.01, \*\*\**p* < 0.001, and \*\*\*\**p* < 0.0001. **e**, Amelioration of cardiac functions in TAC hearts by the RR compounds, which were assessed by echocardiography. Representative images of left ventricular M-mode echocardiographic tracings (left) and results for FS and LV-EDD (right). Data are presented as mean ± SEM (*n* = 9 animals per group). Two-way ANOVA followed by Bonferroni multiple comparison tests were performed. \**p* < 0.05, \*\**p* < 0.01, \*\*\**p* < 0.001, and \*\*\*\**p* < 0.0001. **f**, Amelioration of cardiac functions in TAC hearts by the RR compounds, which were assessed by PV loop analysis. Representative images of PV loops from invasive hemodynamic analysis (upper) and results for dP/dtmax, Tau, Ea, and peak LVP (lower). Data are presented as mean ± SEM (*n* = 5 animals per group). Two-way ANOVA followed by Bonferroni multiple comparison tests were performed. \**p* < 0.05, \*\**p* < 0.01, \*\*\**p* < 0.001, and \*\*\*\**p* < 0.0001.

### Genetic intervention of the pathology

To further validate the concept beyond pharmacological interventions, we utilized a mouse model in which wild-type GAPDH is replaced with a mutant GAPDH [replacement of lysine-225 (K225) with alanine (A)] that is unable to bind with its nuclear chaperon Siah1. With K225 mutation, GAPDH even with C150 posttranslational modification becomes incapable of interacting with Siah1, inhibiting stress-induced nuclear translocation or N-GAPDH cascade by the C150 modification-associated mechanism^13^. In the present study, we generated mice in which this mutant GAPDH was selectively expressed in cardiomyocytes in the heart (GAPDH-K225A mice, K225A) using the tamoxifen-inducible Cre-loxP system, and compared them to mice with wild-type GAPDH (wild-type GAPDH mice, WT) (**Fig. 4a,b, Extended Data Fig. 2, Extended Data Fig. 3a**). The level of *Gapdh* mRNA expression and enzymatic activity were not different between K225A hearts and WT hearts (**Fig. 4c**). Under stress conditions of pressure-overload (TAC), GAPDH-Siah1 binding was evident in WT hearts but was abolished in K225A hearts as expected (**Extended Data Fig. 3b**). Consistent with this biochemical observation, nuclear GAPDH was undetected in K225A TAC hearts, whereas nuclear GAPDH was evident in WT TAC hearts (**Fig. 4d, Extended Data Fig. 3c,d**). These findings confirm that the activation of N-GAPDH cascade elicited by TAC is mediated by the C150 posttranslational modification, GAPDH-Siah1 interaction-mediated mechanism, which is consistent with the pharmacological intervention data described above.

**Fig. 4:**
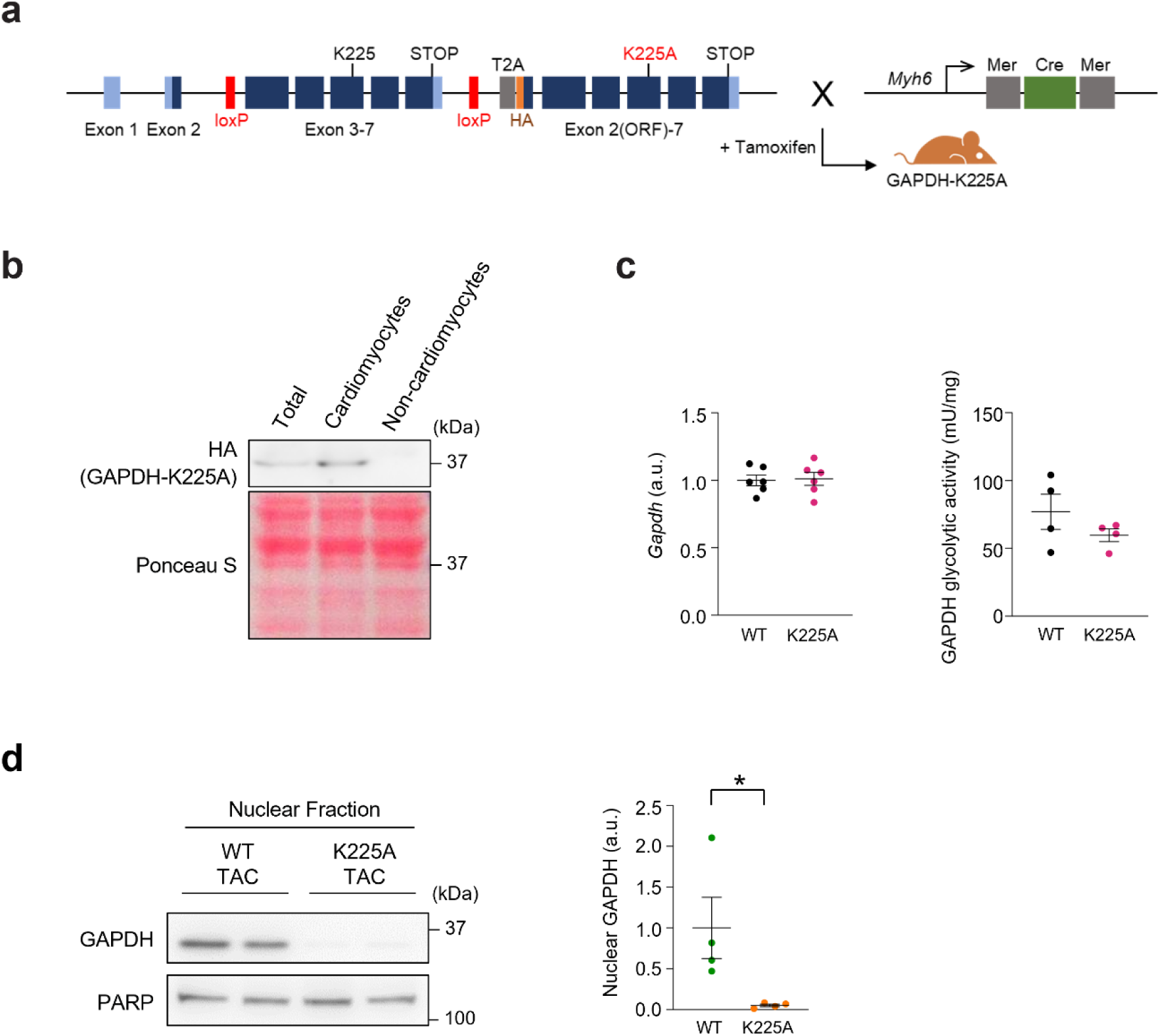
The N-GAPDH cascade is specifically inactivated in cardiomyocytes in mutant (GAPDH-K225A) mice. **a**, A schematic figure of the generation of cardiomyocyte-specific GAPDH-K225A knock-in mice. **b**, Cardiomyocyte-specific introduction of HA-tagged GAPDH-K225A. GAPDH-K225A-HA was expressed in the total heart tissue and enriched in purified cardiomyocytes, but not expressed in non-cardiomyocytes, which was probed with an anti-HA antibody. Ponceau S staining was used as a loading control. **c**, The level of *Gapdh* mRNA expression and enzymatic activity were not different in hearts between mice with wild-type GAPDH (WT) and mice with GAPDH-K225A (K225A). Data are presented as mean ± SEM (*n* = 6 and 4 animals per group for *Gapdh* mRNA and GAPDH glycolytic activity, respectively). Unpaired two-tailed Student’s *t* tests were performed. **d**, Lack of nuclear GAPDH in K225A TAC hearts assessed by immunoblot using an antibody against GAPDH (left) and the quantification results (right). PARP for a loading control of nuclear proteins. Data are presented as mean ± SEM (*n* = 4 animals per group). Unpaired two-tailed Student’s *t* test was performed. \**p* < 0.05.

K225A mice showed a markedly better heart condition compared with WT mice after TAC, as assessed by heart weight, cardiomyocyte size, and fibrosis area (**Fig. 5a,b**). Re-introduction of *Myh7* and other hypertrophy-related molecular changes were almost completely blocked in K225A TAC hearts (**Fig. 5c**). Cardiac function (FS) and chamber remodelling (LV-EDD) by echocardiography were also significantly improved in K225A TAC hearts (**Fig. 5d**). Detailed cardiac functional analyses from PV loop studies further revealed that systolic function (dP/dtmax) was enhanced and diastolic function (Tau) was preserved despite the elevation of chronic cardiac afterload (50% increase in Ea and peak LVP) in K225A TAC hearts, whereas all these functional outcomes were impaired in WT TAC hearts (**Fig. 5e**). These results further support the pivotal role of the N-GAPDH cascade in pressure-overload heart hypertrophy, consistent with the pharmacological data with the RR compound. These molecular intervention data in mice clearly indicate that C150 posttranslational modification, GAPDH-Siah1 interaction-mediated mechanism for stress-induced nuclear GAPDH, plays a crucial role in pressure-overload heart hypertrophy.

**Fig. 5:**
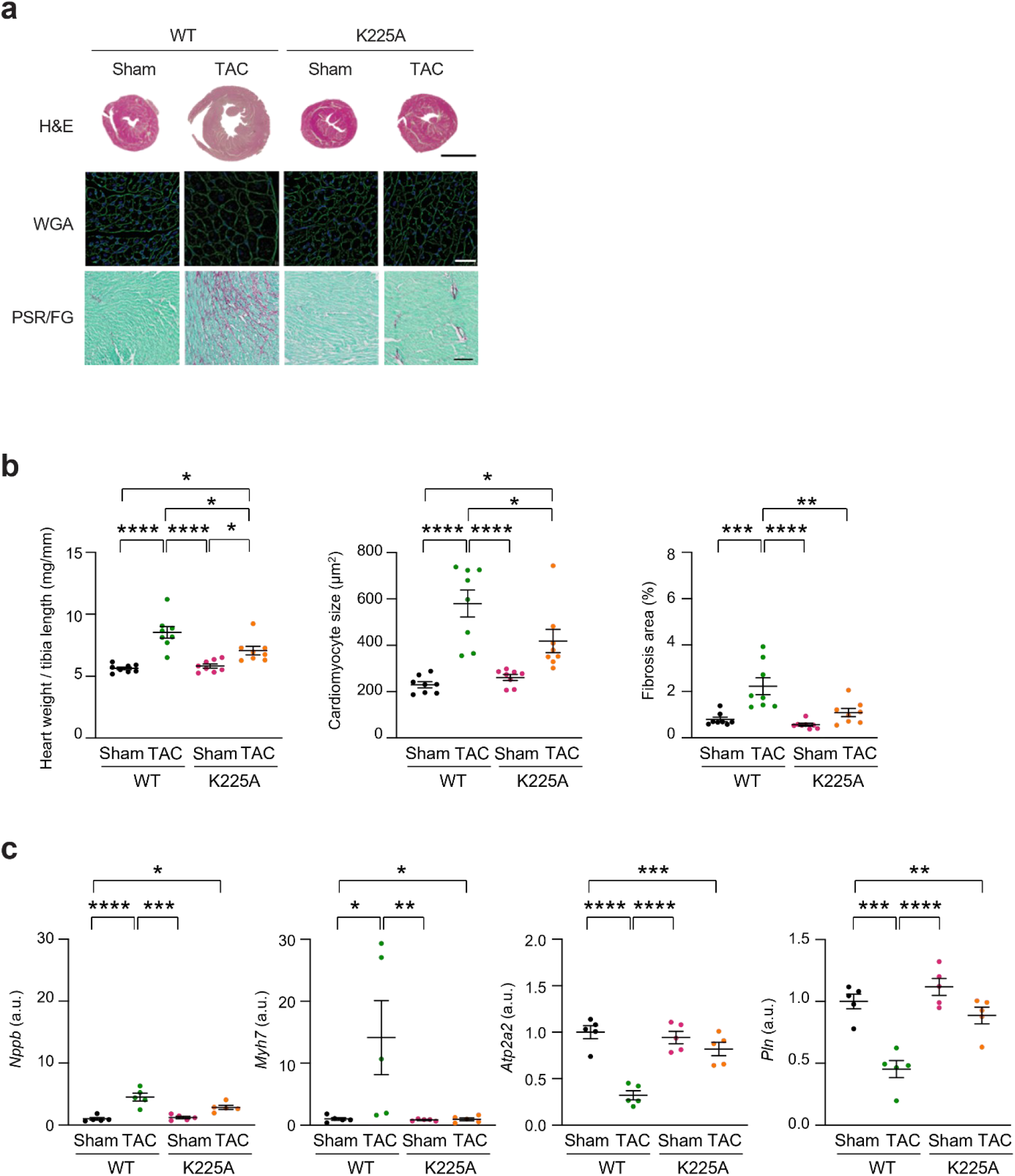
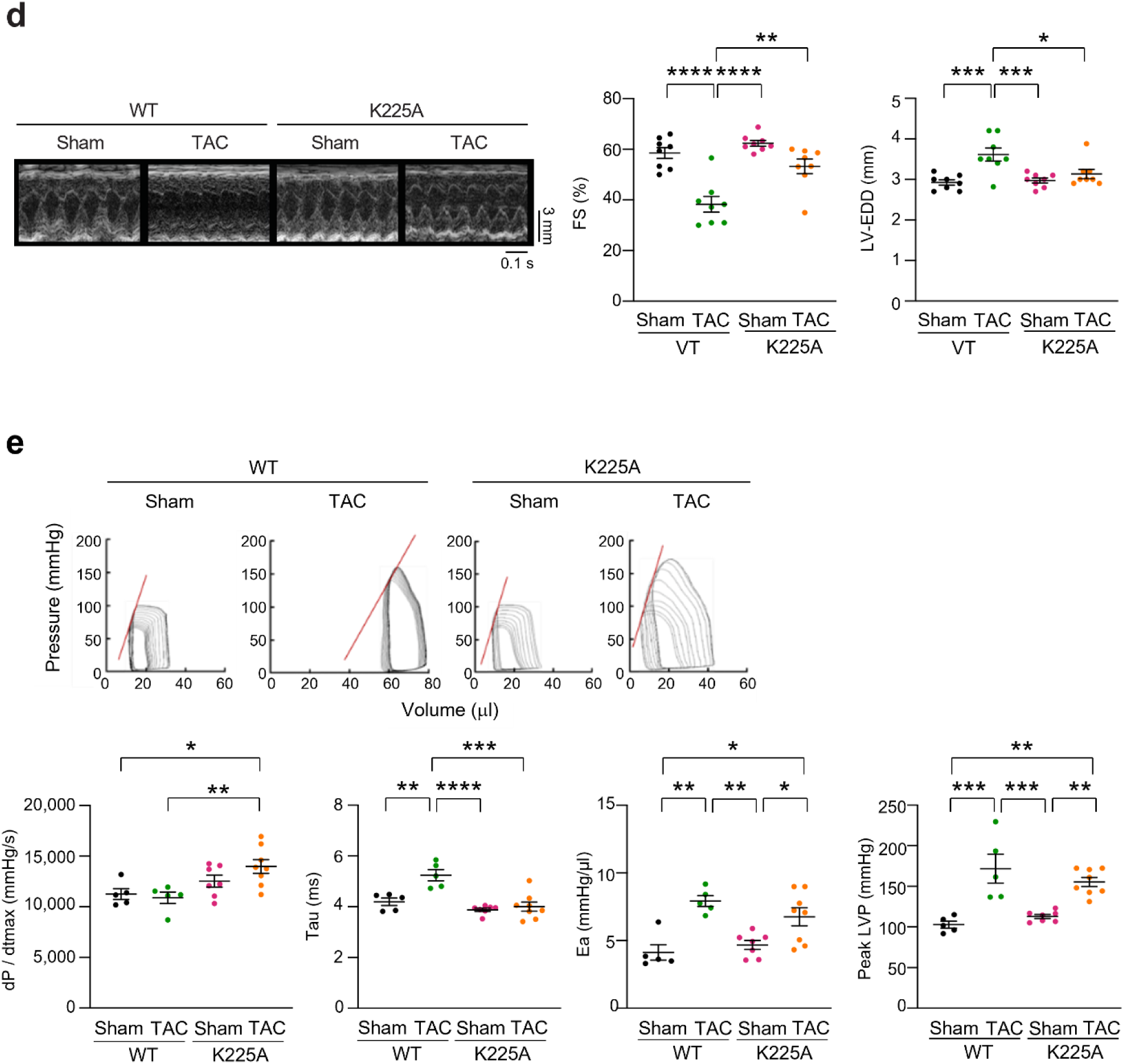
Genetic inactivation of N-GAPDH attenuates heart hypertrophy in mouse TAC model. **a**, Heart hypertrophy with fibrosis elicited by TAC in mice with wild-type GAPDH (WT), which was prevented by genetic inactivation of N-GAPDH in GAPDH-K225A mice (K225A). Representative images of H&E, WGA, and PSR/FG staining of heart cross-sections are shown. Scale bars, 3 mm (H&E); 50 µm (WGA); and 100 µm (PSR/FG). **b**, Heart hypertrophy and fibrosis elicited by TAC, which was prevented by genetic inactivation of N-GAPDH. Heart weight normalized to tibia length, cardiomyocyte size measured from WGA-stained sections, and fibrosis area assessed using PSR/FG staining are shown. Data are presented as mean ± SEM (*n* = 8 animals per group). Two-way ANOVA followed by Bonferroni multiple comparison tests were performed. \**p* < 0.05, \*\**p* < 0.01, \*\*\**p* < 0.001, and \*\*\*\**p* < 0.0001. **c**, Pathological myocardial mRNA expression profiles for *Nppb*, *Myh7*, *Atp2a2,* and *Pln* elicited by TAC, which were prevented by genetic inactivation of N-GAPDH. Data are presented as mean ± SEM (*n* = 5 animals per group). Two-way ANOVA followed by Bonferroni multiple comparison tests were performed. \**p* < 0.05, \*\**p* < 0.01, \*\*\**p* < 0.001, and \*\*\*\**p* < 0.0001. **d**, Cardiac dysfunctions elicited by TAC, which were prevented by genetic inactivation of N-GAPDH. These were assessed by echocardiography. Representative images of left ventricular M-mode echocardiographic tracings (left) and results for FS and LV-EDD (right). Data are presented as mean ± SEM (*n* = 8 animals per group). Two-way ANOVA followed by Bonferroni multiple comparison tests were performed. \**p* < 0.05, \*\**p* < 0.01, \*\*\**p* < 0.001, and \*\*\*\**p* < 0.0001. **e**, Cardiac dysfunctions elicited by TAC, which were prevented by genetic inactivation of N-GAPDH. These were assessed by PV loop analysis. Representative images of PV loops from invasive hemodynamic analysis (upper) and the results for dP/dtmax, Tau, Ea, and peak LVP (lower). Data are presented as mean ± SEM (*n* = 5, 5, 7, and 8 animals for WT-sham, WT-TAC, K225A-sham, and K225A-TAC, respectively). Two-way ANOVA followed by Bonferroni multiple comparison tests were performed. \**p* < 0.05, \*\**p* < 0.01, \*\*\**p* < 0.001, and \*\*\*\**p* < 0.0001.

### N-GAPDH, BRG1/HDAC2, and the *Myh* isoform shift

To elucidate the mechanism of heart hypertrophy mediated by nuclear-translocated, pathological GAPDH, we performed an unbiased proteomic analysis, looking for proteins associated with GAPDH in the nucleus of TAC hearts. Nuclear proteins from TAC hearts were immunoprecipitated using a GAPDH antibody or control IgG, followed by liquid chromatography-tandem mass spectrometry (LC-MS/MS) analysis (**Fig. 6a**). This unbiased, comprehensive analysis identified 567 proteins as potential interactors of nuclear GAPDH (**Fig. 6b**). Gene Ontology (GO) analysis showed that half of the top ten pathways (terms), ranked by the number of identified proteins (**Fig. 6c**), were related to transcriptional regulation and chromatin remodelling^32,33^. In particular, we found multiple molecules of the Switch/Sucrose Non-fermentable (SWI/SNF) complex, which are essential for chromatin remodelling and histone modification, as candidate proteins that interact with nuclear GAPDH (**Supplementary Table 3**). Among these candidates, we paid a particular attention to two molecules: BRG1 and HDAC2. BRG1 is an ATPase subunit of the SWI/SNF complex, which plays a crucial role in regulating the isoform shift on *Myh* (from adult-type *Myh6* to its fetal form *Myh7*) as well as *Myh7*-coded lncRNA [*Myosin-heavy-chain-associated RNA transcript (Mhrt)*] for hypertrophy development. Since BRG1 needs to interact with HDAC2 to activate this process^27,28^, we assessed the role of nuclear GAPDH as a molecular scaffold to facilitate the BRG1-HDAC2 interaction.

**Fig. 6:**
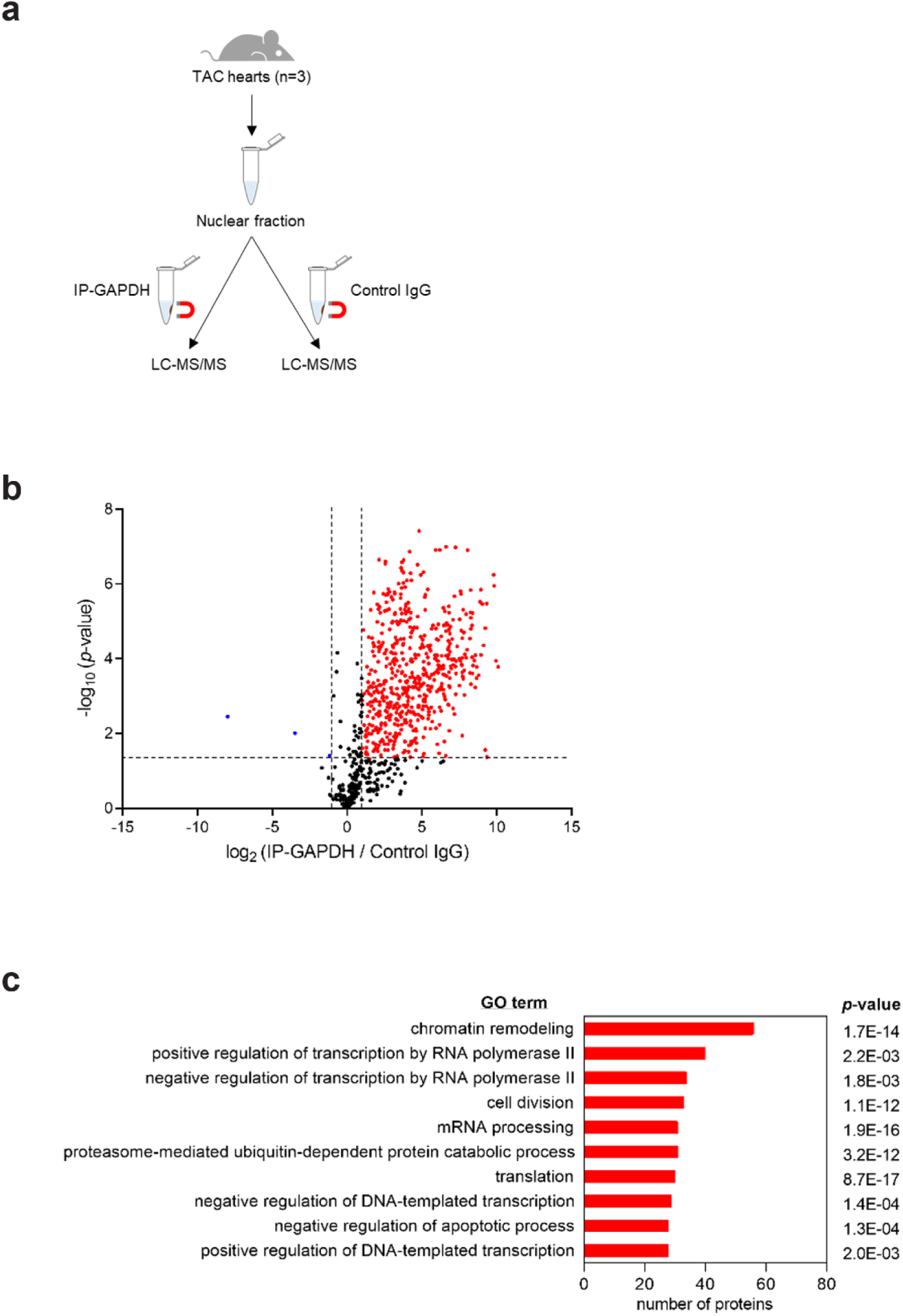
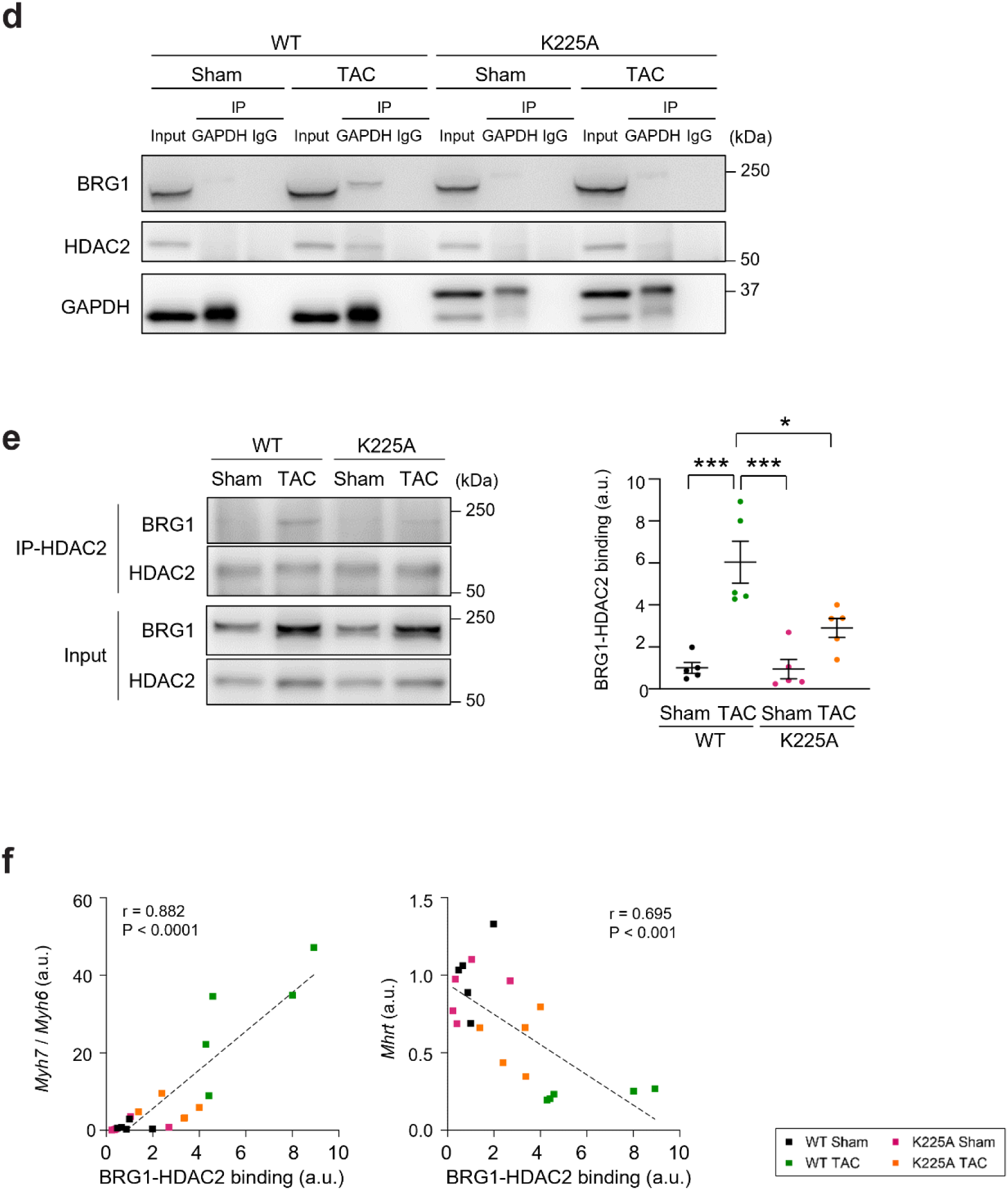
N-GAPDH enhances BRG1-HDAC2 interaction to cause heart hypertrophy. **a**, Schematic representation of the proteomic analysis. Nuclear fractions of TAC hearts in C57BL/6J mice (*n* = 3) were immunoprecipitated with an anti-GAPDH antibody (IP-GAPDH) or control IgG, followed by LC-MS/MS. **b**, Volcano plot of proteins identified by LC-MS/MS comparing the IP-GAPDH and control IgG groups. A total of 567 proteins significantly enriched in the IP-GAPDH group (fold change > 2, *p* < 0.05) are highlighted in red. **c**, GO analysis of the 567 proteins enriched in the IP-GAPDH group. The top 10 GO terms in biological processes are ranked based on the number of associated proteins. **d**, Interactions of GAPDH with BRG1 and HDAC2 by co-immunoprecipitation from heart lysates. GAPDH-BRG1 interaction and GAPDH-HDAC2 interaction were induced by TAC in wild-type GAPDH (WT) hearts but not in GAPDH-K225A (K225A) hearts. The upper bands in the GAPDH panel correspond mutant GAPDH-K225A which is HA-tagged. **e**, Interaction between BRG1 and HDAC2 by co-immunoprecipitation from heart lysates (left). BRG1-HDAC2 interaction was robustly induced by TAC in WT but only weakly in K225A hearts. Quantification results (right) are presented as mean ± SEM (*n* = 5 animals per group). Two-way ANOVA followed by Bonferroni multiple comparison tests were performed. \**p* < 0.05 and \*\*\**p* < 0.001. **f**, Correlations between BRG1-HDAC2 binding and the *Myh7*/*Myh6* ratio (left) or *Mhrt* expression (right) (*n* = 5 animals per group), with Pearson correlation coefficient (*r*), *p*-values, and fitted trend lines.

We first confirmed the interactions between GAPDH and BRG1, as well as GAPDH and HDAC2 biochemically. Given that BRG1 is reportedly upregulated by TAC^27^, we utilized IgG as a negative control for co-immunoprecipitation, and assessed the specific binding between GAPDH and BRG1. In WT TAC hearts, both BRG1 and HDAC2 co-immunoprecipitated with GAPDH, whereas neither interaction was detected in WT sham hearts (**Fig. 6d, Extended Data Fig. 4**). By contrast, these interactions were barely detected in K225A TAC hearts, as well as K225A sham hearts (**Fig. 6d, Extended Data Fig. 4**). Next, we assessed the contribution of nuclear GAPDH to the binding of BRG1 with HDAC2, which induces *Myh7*^27,28^. In WT hearts, BRG1-HDAC2 binding was markedly induced by TAC^27^, whereas this interaction was only weakly induced by TAC in K225A hearts (**Fig. 6e**). These findings indicate that nuclear-translocated GAPDH serves as a scaffold to enhance the BRG1-HDAC2 interaction, or the requirement of the N-GAPDH cascade for the BRG1-HDAC2 interaction, in response to extrinsic stress of pressure-overload.

Considering that the BRG1-HDAC2 protein interaction mediates the fundamental process for heart hypertrophy via the *Myh7/Myh6* ratio shift^27,28^, we quantitatively addressed the relationship between the levels of BRG1-HDAC2 binding and the *Myh7/Myh6* ratio shift. We also quantified the expression of *Mhrt* whose reduction mediated by the BRG1-HDAC2 interaction further facilitates the *Myh7/Myh6* ratio shift^28^. As shown in **Fig. 6f**, the levels of BRG1-HDAC2 protein interaction significantly paralleled the *Myh7/Myh6* ratio. Consistent with this, a significantly negative correlation was also observed between the protein interaction and *Mhrt* expression (**Fig. 6f**).

Collectively, these findings support the pivotal role for the N-GAPDH cascade serving as a molecular scaffold to enhance BRG1-HDAC2 binding, thereby contributing to heart hypertrophy via the regulation of specific gene expressions.

## Discussion

In the present study, we demonstrated that stress-induced GAPDH nuclear translocation (N-GAPDH cascade) is a causal role in heart hypertrophy *in vivo*. We provided evidence that supported the notion at multiple levels, from the molecular to cardiac physiological levels, with two independent methods (genetic and pharmacological interventions). In the pharmacological intervention, we used the RR compound, which is a structural analogue of a clinically available medicine Deprenyl. Accordingly, RR is supposed to have a safety profile, which supports the translational potential of targeting N-GAPDH in heart disease.

Previous studies have outlined a framework in which cellular stress triggers posttranslational modifications of GAPDH, leading to nuclear translocation, where it may recruit and regulate histone acetyltransferase (HAT) / HDAC-associated molecules in the nucleus. Nevertheless, the direct link between the regulation of GAPDH on its nuclear protein interactors and specific gene transcription for the final phenotype has been unclear. In the present work, we demonstrate that the N-GAPDH cascade directly mediates the stress-elicited regulation of the BRG1-HDAC2 interaction, critically affecting specific gene expression (the *Myh7/Myh6* ratio shift and *Mhrt* expression) for pathological heart hypertrophy. We believe that the present collective findings provide strong support for the overall concept of the N-GAPDH cascade, a stress-induced signal transduction that regulates HAT / HDAC-associated molecules and gene expression for a stress-elicited final cellular phenotype.

We believe that the present work may impact a fundamental biological framework beyond its medical implications. Stress-induced reversion of a cellular phenotype from adult to immature or fetal state has been frequently seen in living organisms^34–36^. Tumorigenesis in adulthood is also interpreted in this context^37^. Nonetheless, it was unclear which specific mechanism(s) mediates the stress-induced reversion. Here we provide evidence that the N-GAPDH cascade plays a role in this evolutionally conserved and fundamental phenotype in living organisms. Given that GAPDH is an evolutionally conserved molecule, its novel role for the fundamental mechanism shown in the present study may be biologically reasonable. Collectively, we provide evidence for the central role of GAPDH in cellular homeostatic control, serving as a sensor to diverse stress and regulating fundamental cellular response.

## Acknowledgments

We thank Dr. Pamela Talalay and Ms. Lauren Guttman for critical reading of the manuscript. We also thank Ms. Yukiko Lema for organizing the figures and manuscript. This work was supported by USPHS grants of HL-093432 (E.T.), MH-084018 (A.S.), MH-094268 (A.S.), MH-069853 (A.S.), MH-085226 (A.S.), MH-088753 (A.S.), MH-092443 (A.S.), MH-105660 (A.S.), MH-107730 (A.S.), HL-077180 (D.A.K.), HL-059408 (D.A.K.), HL-07227 (M.Z.), and F31NS070459 (C.T.), AHA grant GIA 7700071 (E.T.), Miyata Cardiac Research Promotion Foundation grant (E.T.), JSPS KAKENHI grants of 24K02441 (E.T.), 21K08048 (E.T.), 24K19053 (S.N.), 22KJ1202 (S.N.), and 22J10789 (S.N.), Fukuda Foundation for Medical Technology (E.T., G.N., S.N.), Japan Heart Foundation research grant (G.N.), American Heart Association post-doctoral fellowship (M.Z.), Stanley (A.S.), RUSK/S-R (A.S.), NARSAD (A.S., H.J-P., and N.S.), and Maryland Stem Cell Research Fund (A.S.).

## Author contributions

E.T. and A.S. conceptualized all the project. S.N., G.N., M.Z., T.K., H.S., M.S., A.R., N.K., G.Z., M.I., T.T., D.L., C.T., N.S., Y.T., H.J.-P., T. Saitoh and K.I. performed all experiments and analyses. E.A., M.H., M.A., Y.Y., T.U., and M.O. contributed to the acquisition, processing, and interpretation of clinical samples. E.T. and A.S. wrote the manuscript. B.S., T. Sasano, and D.A.K. provided critical edits and feedback to the manuscript.

## Supplementary information

Supplementary information is available for this paper.

## References

1 Sirover, M. A. Moonlighting glyceraldehyde-3-phosphate dehydrogenase: posttranslational modification, protein and nucleic acid interactions in normal cells and in human pathology. Crit Rev Biochem Mol Biol 55, 354–371 (2020).

2 Tristan, C., Shahani, N., Sedlak, T. W. & Sawa, A. The diverse functions of GAPDH: views from different subcellular compartments. Cellular signalling 23, 317–323 (2011).

3 Sirover, M. A. On the functional diversity of glyceraldehyde-3-phosphate dehydrogenase: biochemical mechanisms and regulatory control. Biochim Biophys Acta 1810, 741–751 (2011).

4 Chuang, D. M., Hough, C. & Senatorov, V. V. Glyceraldehyde-3-phosphate dehydrogenase, apoptosis, and neurodegenerative diseases. Annu Rev Pharmacol Toxicol 45, 269–290 (2005).

5 Shahani, N. & Sawa, A. Nitric oxide signaling and nitrosative stress in neurons: role for S-nitrosylation. Antioxid Redox Signal 14, 1493–1504 (2011).

6 Hara, M. R., Cascio, M. B. & Sawa, A. GAPDH as a sensor of NO stress. Biochim Biophys Acta 1762, 502–509 (2006).

7 Shin, M. K. et al. Reducing acetylated tau is neuroprotective in brain injury. Cell 184, 2715–2732.e2723 (2021).

8 Yoon, S. et al. S-Nitrosylation of Histone Deacetylase 2 by Neuronal Nitric Oxide Synthase as a Mechanism of Diastolic Dysfunction. Circulation 143, 1912–1925 (2021).

9 Sirover, M. A. Subcellular dynamics of multifunctional protein regulation: mechanisms of GAPDH intracellular translocation. J Cell Biochem 113, 2193–2200 (2012).

10 Sirover, M. A. New nuclear functions of the glycolytic protein, glyceraldehyde-3-phosphate dehydrogenase, in mammalian cells. J Cell Biochem 95, 45–52 (2005).

11 Galván-Peña, S. et al. Malonylation of GAPDH is an inflammatory signal in macrophages. Nature communications 10, 338 (2019).

12 Chang, C. H. et al. Posttranscriptional control of T cell effector function by aerobic glycolysis. Cell 153, 1239–1251 (2013).

13 Hara, M. R. et al. S-nitrosylated GAPDH initiates apoptotic cell death by nuclear translocation following Siah1 binding. Nat Cell Biol 7, 665–674 (2005).

14 Singh, R. & Green, M. R. Sequence-specific binding of transfer RNA by glyceraldehyde-3-phosphate dehydrogenase. Science 259, 365–368 (1993).

15 Morgenegg, G. et al. Glyceraldehyde-3-phosphate dehydrogenase is a nonhistone protein and a possible activator of transcription in neurons. J Neurochem 47, 54–62 (1986).

16 Sirover, M. A. Pleiotropic effects of moonlighting glyceraldehyde-3-phosphate dehydrogenase (GAPDH) in cancer progression, invasiveness, and metastases. Cancer Metastasis Rev 37, 665–676 (2018).

17 Boradia, V. M., Raje, M. & Raje, C. I. Protein moonlighting in iron metabolism: glyceraldehyde-3-phosphate dehydrogenase (GAPDH). Biochem Soc Trans 42, 1796–1801 (2014).

18 Sawa, A., Khan, A. A., Hester, L. D. & Snyder, S. H. Glyceraldehyde-3-phosphate dehydrogenase: nuclear translocation participates in neuronal and nonneuronal cell death. Proc Natl Acad Sci U S A 94, 11669–11674 (1997).

19 Sen, N. et al. Nitric oxide-induced nuclear GAPDH activates p300/CBP and mediates apoptosis. Nat Cell Biol 10, 866–873 (2008).

20 Chang, C. et al. AMPK-Dependent Phosphorylation of GAPDH Triggers Sirt1 Activation and Is Necessary for Autophagy upon Glucose Starvation. Mol Cell 60, 930–940 (2015).

21 Martin, T. G., Juarros, M. A. & Leinwand, L. A. Regression of cardiac hypertrophy in health and disease: mechanisms and therapeutic potential. Nature reviews. Cardiology 20, 347–363 (2023).

22 Takimoto, E. et al. Chronic inhibition of cyclic GMP phosphodiesterase 5A prevents and reverses cardiac hypertrophy. Nat.Med. 11, 214–222 (2005).

23 Takimoto, E. et al. Regulator of G protein signaling 2 mediates cardiac compensation to pressure overload and antihypertrophic effects of PDE5 inhibition in mice. J Clin Invest 119, 408–420 (2009).

24 Zhang, M. et al. Myocardial remodeling is controlled by myocyte-targeted gene regulation of phosphodiesterase type 5. Journal of the American College of Cardiology 56, 2021–2030 (2010).

25 Dorn, G. W., 2nd & Brown, J. H. Gq signaling in cardiac adaptation and maladaptation. Trends in cardiovascular medicine 9, 26–34 (1999).

26 Dorn, G. W., 2nd & Force, T. Protein kinase cascades in the regulation of cardiac hypertrophy. J Clin Invest 115, 527–537 (2005).

27 Hang, C. T. et al. Chromatin regulation by Brg1 underlies heart muscle development and disease. Nature 466, 62–67 (2010).

28 Han, P. et al. A long noncoding RNA protects the heart from pathological hypertrophy. Nature 514, 102–106 (2014).

29 Li, P., Ge, J. & Li, H. Lysine acetyltransferases and lysine deacetylases as targets for cardiovascular disease. Nature reviews. Cardiology 17, 96–115 (2020).

30 Ramos, A. et al. Nuclear GAPDH in cortical microglia mediates cellular stress-induced cognitive inflexibility. Mol Psychiatry 10, 2967–2978 (2024).

31 Wettschureck, N. et al. Absence of pressure overload induced myocardial hypertrophy after conditional inactivation of Galphaq/Galpha11 in cardiomyocytes. Nat Med 7, 1236–1240 (2001).

32 Clapier, C. R., Iwasa, J., Cairns, B. R. & Peterson, C. L. Mechanisms of action and regulation of ATP-dependent chromatin-remodelling complexes. Nat Rev Mol Cell Biol 18, 407–422 (2017).

33 Suganuma, T. & Workman, J. L. Signals and combinatorial functions of histone modifications. Annu Rev Biochem 80, 473–499 (2011).

34 Waheed, Z. et al. The Role of Tau Proteoforms in Health and Disease. Mol Neurobiol 60, 5155–5166 (2023).

35 Chanda, D. et al. Developmental pathways in the pathogenesis of lung fibrosis. Mol Aspects Med 65, 56–69 (2019).

36 Sakamoto, T. & Kelly, D. P. Cardiac maturation. J Mol Cell Cardiol 187, 38–50 (2024).

37 Fey, S. K., Vaquero-Siguero, N. & Jackstadt, R. Dark force rising: Reawakening and targeting of fetal-like stem cells in colorectal cancer. Cell Rep 43, 114270 (2024).

