## Supplementary material for "Nuclear GAPDH cascade mediates stress-dependent pathological cardiac growth": Supplmentary Materials

#### Materials and methods

##### Animal models

Male mice (8-12 weeks old) were used for all experiments, with female mice included in a subset of experiments. C57BL/6J mice were purchased from Jackson Laboratory. Conditional knock-in mice were generated on a C57BL/6J background by UNITECH (Chiba, Japan) (see **Extended Data Fig. 2a**). A floxed mouse line was established in which a single mutation was introduced into exon 5 of the *Gapdh* gene to replace lysine-225 with alanine (K225A). $Gapdh^{K225A\ fl/fl}$ mice were crossed with *Myh6*-MerCreMer transgenic mice (Jackson Laboratory) to generate mice with tamoxifen-inducible cardiomyocyte-specific GAPDH-K225A ($Gapdh^{K225A\ fl/fl}$/*Myh6*-MerCreMer). Tamoxifen (20 mg/kg/day i.p.) was administered for three days[38] to induce Cre-mediated recombination for cardiomyocyte-specific expression of GAPDH-K225A (K225A mice). $Gapdh^{K225A\ fl/fl}$ mice also received tamoxifen, serving as controls (mice with wild-type GAPDH, WT). TAC surgery inducing cardiac pressure overload[22-24] was performed four weeks later to allow maximum recombination as well as recovery from tamoxifen-induced heart damage[39,40]. This tamoxifen treatment does not interfere with the experiments that examine the N-GAPDH cascade, whereas it slightly affects molecular results regarding the TAC response. All experiments in TAC-operated mice were performed 10 days after surgery. Echocardiography and PV loop analysis were performed as previously described[22-24]. All procedures were approved by the Animal Care and Use Committees of Johns Hopkins University and The University of Tokyo.

##### Cell cultures

H9c2 cells were seeded and stimulated the next day with 0.05 μM ET-1, 5 μM angiotensin II, or 100 μM phenylephrine for 48 h. Neonatal rat cardiomyocytes were isolated from 1- to 2-day-old Sprague-Dawley rats and stimulated with 0.05 μM ET-1[22], with or without 1 nM RR compound for 48 h. Protein synthesis was assessed by $^3$H-leucine incorporation[22,23].

##### RR compound

The synthesis and characterization of RR compound were reported previously[30,41]. Off-target activity was assessed using the HitProfilingScreen® radiolabelled binding assay (Eurofins Panlab Inc.) at a concentration of 1 μM.

#### Isolation of cardiac cells

Cardiac cells were isolated using the Langendorff perfusion technique. After euthanasia, the heart was excised, cannulated through the aorta, and retrogradely perfused at 37 °C for 2 min with cell isolation buffer (25 mM HEPES, pH 7.4, 130 mM NaCl, 5.4 mM KCl, 0.5 mM $MgCl_2$, 0.33 mM $NaH_2PO_4$, 22 mM D-glucose) supplemented with 0.4 mM EGTA. Subsequently, the heart was perfused at 37 °C for 10 min with the cell isolation buffer supplemented with 0.1 mM $CaCl_2$, 1 mg/mL collagenase type II (Worthington), and 0.05 mg/mL protease XIV (Sigma). The heart was then dissociated into single cells in the buffer supplemented with an additional 2 mg/mL bovine serum albumin. The suspension was centrifuged at 300 rpm for 2 min to obtain the cardiomyocyte fraction, and the supernatant was further centrifuged at 3000 rpm for 5 min to obtain the non-cardiomyocyte fraction.

#### Histological analysis

Heart samples were fixed with 10% formalin, embedded in paraffin, sectioned, and stained for cardiomyocyte size and fibrosis as previously described[24,42]. For immunofluorescence, tissue sections were stained with antibodies against GAPDH (Cell Signaling Technology, 2118; Proteintech, 60004-1-Ig), cardiac troponin T (Proteintech, 15513-1-AP), actin (Cell Signaling Technology, 4968), or HA-tag (Roche, 11867423001). Isolated cardiomyocytes were fixed with a 1:1 methanol-acetone mixture and stained with antibodies against GAPDH or sulphonated GAPDH (Cys150) as previously described[23,24,30]. Histological images were acquired using a fluorescence microscope (APX100, Evident) and a confocal laser microscope (FV3000, Evident).

#### Protein analysis

Total lysates of cells and tissues were prepared using RIPA lysis buffer supplemented with a protease inhibitor cocktail (Roche). Subcellular fractionation into cytosolic and nuclear soluble fractions was performed using the LysoPure Nuclear and Cytoplasmic Extractor Kit (FUJIFILM Wako Pure Chemical Corporation), following the manufacturer's instructions. Co-immunoprecipitation for the GAPDH-Siah1 protein interaction was performed as previously described[30]. For co-immunoprecipitation of GAPDH and HDAC2, heart tissues were homogenized in NP-40 lysis buffer (25 mM HEPES, pH 7.5, 250 mM KCl, 12.5 mM $MgCl_2$, 0.5% NP-40, 8% glycerol, 1 mM dithiothreitol, and a protease inhibitor cocktail). Lysates were diluted with 2 volumes of lysis buffer containing 100 mM KCl and incubated with an anti-GAPDH antibody (Abcam, ab181602), an anti-HDAC2 antibody (Abcam, ab16032), or an isotype control IgG, with rotation at 4 °C, overnight. Protein G magnetic beads were then added and incubated for 2 h at 4 °C with rotation. Beads were washed twice with lysis buffer containing 150 mM KCl, and bound protein complexes were eluted for further analysis. Protein extracts were analysed by Western blotting as previously described[22-24]. The following antibodies were used: anti-GAPDH (Abcam, ab181602), anti-LMNB1 (Cell Signaling Technology, 13435), anti-LDH (Cell Signaling Technology, 2012), anti-HA-tag (Santa Cruz Biotechnology, sc-57592), anti-Siah1 (Santa Cruz Biotechnology, sc-5505 and sc-5506), anti-PARP (Cell Signaling Technology, 9532), anti-BRG1 (Abcam, ab110641), and anti-HDAC2 (Cell Signaling Technology, 57156).

**Proteomic analysis**

Nuclear proteins were extracted from the left ventricular tissues of TAC mice (C57BL/6J) using the LysoPure Nuclear and Cytoplasmic Extractor Kit (FUJIFILM Wako Pure Chemical Corporation), following the manufacturer's instructions. Protein G magnetic beads were pre-incubated with either an anti-GAPDH antibody (Abcam, ab181602) or isotype control IgG, then added to the nuclear extracts and incubated for 1 h at room temperature. The immunoprecipitated proteins were analysed by LC-MS/MS for protein identification and quantification. LC-MS/MS measurements, including protein quantification, data processing, and statistical analysis, were performed by Kazusa Genome Technologies Inc. (Chiba, Japan). GO enrichment analysis was conducted using the Database for Annotation, Visualization, and Integrated Discovery (DAVID)[43,44].

**RNA analysis**

Total RNA was extracted from cells or heart tissues using TRIzol (Invitrogen) or TRI Reagent (Molecular Research Center), following the manufacturer’s instructions. Reverse transcription was performed using the SuperScript First-Strand Synthesis System (Invitrogen) or ReverTra Ace qPCR RT Master Mix (TOYOBO). Quantitative PCR was performed using TaqMan PCR

Master Mix reagent (Applied Biosystems) or THUNDERBIRD SYBR qPCR Mix (TOYOBO). Gene expression levels were quantified using the $2^{-\Delta\Delta Ct}$ method and normalized to 18S rRNA as an internal control.

**GAPDH activity assay**

Heart tissues were collected and analysed for GAPDH enzymatic activity using the Glyceraldehyde-3-Phosphate Dehydrogenase Activity Assay Kit (Abcam), following the manufacturer's instructions.

**Human studies**

All procedures involving human myocardial samples were approved by the institutional ethics committee of The University of Tokyo (approval number: 2024386NI). Left ventricular myocardial tissues from heart failure patients with dHCM were collected during left ventricular assist device implantation. Control samples were obtained from right ventricular endomyocardial biopsies performed one year after heart transplantation, and only those without histological evidence of transplant rejection were included. Patient characteristics are summarized in **Supplementary Table 1**.

**Statistical analysis**

Statistical analyses were performed using GraphPad Prism version 9.4.0. Data are presented as mean ± SEM. Comparisons between two groups were made using an unpaired two-tailed Student's *t* test. Comparisons among more than two groups were performed using one-way or two-way analysis of variance (ANOVA), followed by Bonferroni multiple comparison test. Pearson's correlation coefficient ($r$) was used for correlation analysis. Statistical significance was defined as $p < 0.05$.

**Data availability**

The mass spectrometry proteomics data have been deposited to the ProteomeXchange Consortium via the PRIDE partner repository with the dataset identifier PXD062010. The dataset is currently under restricted access and will be made publicly available upon journal publication.

**Method references**


38 Deloux, R. *et al.* Voluntary Exercise Improves Cardiac Function and Prevents Cardiac Remodeling in a Mouse Model of Dilated Cardiomyopathy. *Frontiers in physiology* **8**, 899 (2017).

39 Asp, M. L., Martindale, J. J. & Metzger, J. M. Direct, differential effects of tamoxifen, 4-hydroxytamoxifen, and raloxifene on cardiac myocyte contractility and calcium handling. *PLoS One* **8**, e78768 (2013).

40 Meng, T. *et al.* Tamoxifen induced cardiac damage via the IL-6/p-STAT3/PGC-1α pathway. *Int Immunopharmacol* **125**, 110978 (2023).

41 Sawa, A., Takimoto, E., Shahani, N., Kass, D. & Saito, T. GAPDH cascade inhibitor compounds and methods of use and treatment of stress induced disorders including mental illness. United States patent US10292980B2 (2019).

42 Sano, S. *et al.* TP53-mediated therapy-related clonal hematopoiesis contributes to doxorubicin-induced cardiomyopathy by augmenting a neutrophil-mediated cytotoxic response. *JCI Insight* **6** (2021).

43 Sherman, B. T. *et al.* DAVID: a web server for functional enrichment analysis and functional annotation of gene lists (2021 update). *Nucleic Acids Res* **50**, W216–w221 (2022).

44 Huang da, W., Sherman, B. T. & Lempicki, R. A. Systematic and integrative analysis of large gene lists using DAVID bioinformatics resources. *Nat Protoc* **4**, 44–57 (2009).

#### Extended Data Fig. 1

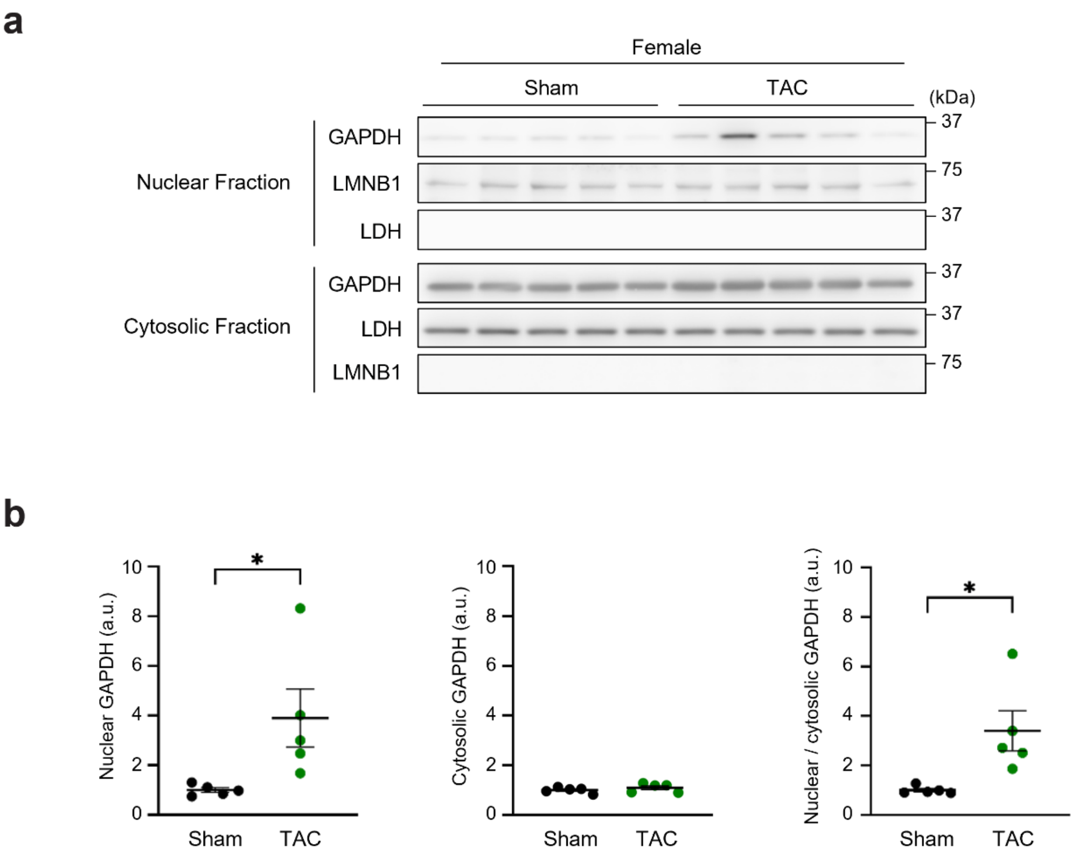

**Extended Data Fig. 1: Nuclear translocation of GAPDH in TAC hearts of female mice.**

**a**, N-GAPDH shown by immunoblots using nuclear and cytosolic fractions from sham or TAC hearts from female mice, with LMNB1 and LDH as loading controls for nuclear and cytosol proteins, respectively.

**b**, Quantification results are presented as mean ± SEM ($n$ = 5 animals per group). Unpaired two-tailed Student's $t$ tests were performed. *$p < 0.05$.

#### Extended Data Fig. 2

**a**

mouse Gapdh (NM_008084.4)

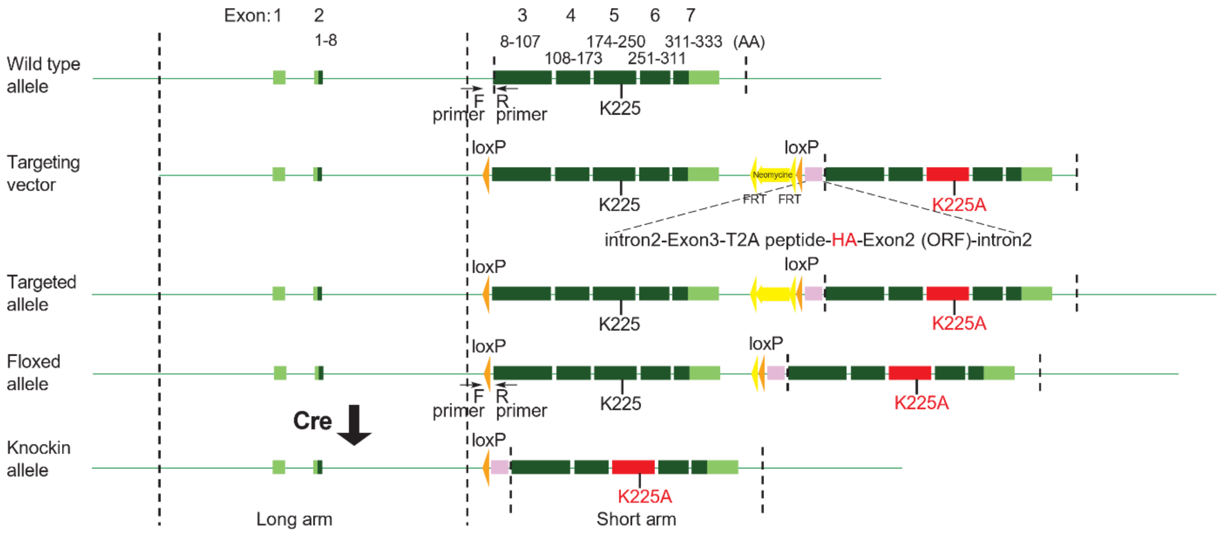


**b**

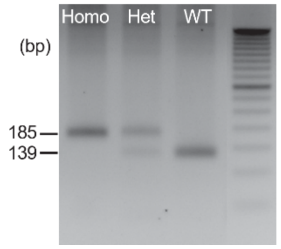


Foward (F) primer: ACTTTCTAGGGTGGGAACAGTCTAT
Reverse (R) primer: TACAAGCCTTTATGCAGACACAGTA

**Extended Data Fig. 2: Generation and genotyping of GAPDH-K225A mice.**

**a**, Schematic representation of the GAPDH-K225A conditional knock-in strategy. The targeting vector was composed of *Gapdh* gene segments, a short gene fragment (short arm) and a larger fragment (long arm). The short arm contains exons 3-7, with the K225A mutation introduced in exon 5. Cre-mediated recombination at loxP sites results in the expression of HA-tagged GAPDH-K225A. The locations of the forward (F) and reverse (R) PCR primers used for genotyping are indicated.

**b**, Genotyping PCR analysis of tail genomic DNA from mice with the following genotypes: $Gapdh^{K225A\ fl/fl}$ (lane 1, Homo), $Gapdh^{K225A\ fl/WT}$ (lane 2, Het), and $Gapdh^{WT/WT}$ (lane 3, WT). The sequences of the primers used are also shown.

### Extended Data Fig. 3

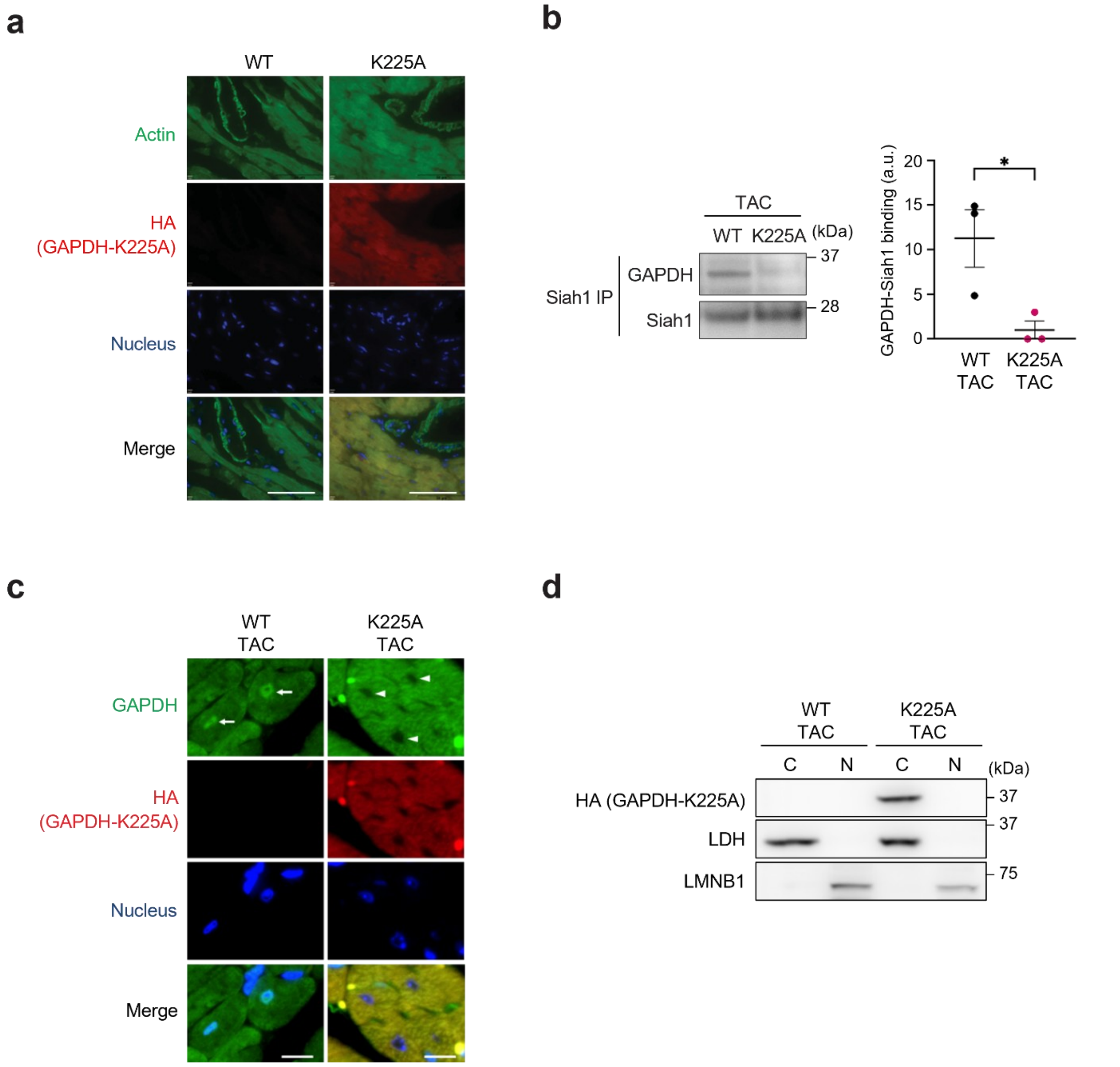

**Extended Data Fig. 3: Abolition of GAPDH-Siah1 binding and nuclear translocation of GAPDH by TAC in GAPDH-K225A mice.**

**a**, Cardiomyocyte-specific introduction of GAPDH-K225A by immunofluorescence imaging. Cardiomyocytes and vascular cells are stained with actin (green). HA-tagged mutant GAPDH-K225A (red) is exclusively expressed in cardiomyocytes (yellow in a Merge panel in K225A) and not in vascular tissues (green in a Merge panel in K225A). Blue, DAPI (nucleus). Scale bar, 50 μm.

**b**, GAPDH-Siah1 binding by co-immunoprecipitation from heart lysates in TAC hearts. GAPDH-Siah1 binding was evident in WT TAC hearts, but was abolished in GAPDH-K225A TAC hearts (left). The quantification results (right). Data are presented as mean ± SEM ($n$ = 3 animals per group). Unpaired two-tailed Student's $t$ test was performed. *$p < 0.05$.

**c**, Nuclear translocation of GAPDH assessed by immunofluorescence imaging. N-GAPDH was not observed in GAPDH-K225A TAC hearts (arrowheads) with an antibody against GAPDH or HA, in contrast to robust nuclear translocation in WT TAC hearts (arrows). Green, GAPDH; red, HA; blue, DAPI (nucleus). Scale bar, 10 μm.

**d**, Abolition of nuclear translocation of GAPDH-K225A. GAPDH-K225A remained in cytosol in TAC hearts by immunoblot using an antibody against HA. C, cytosolic fraction; N, nuclear fraction. LDH and LMNB1 for loading controls of cytosol and nuclear proteins, respectively.

### Extended Data Fig. 4

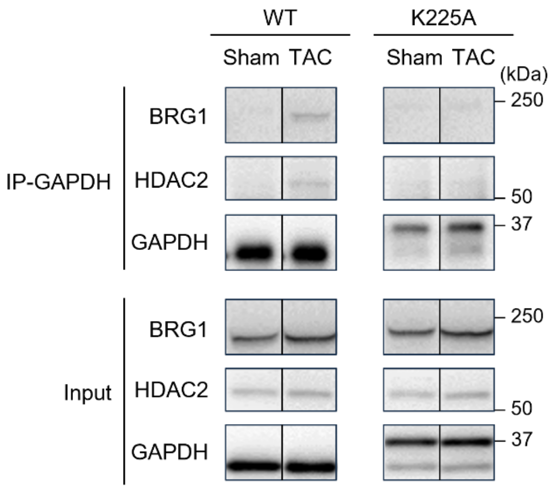

**Extended Data Fig. 4: Abolition of TAC-induced interactions of GAPDH with BRG1 and HDAC2 in GAPDH-K225A mice.**

Interactions of GAPDH with BRG1 and HDAC2 by co-immunoprecipitation from heart lysates. GAPDH-BRG1 interaction and GAPDH-HDAC2 interaction were induced by TAC in WT hearts but not in K225A hearts. The upper bands in the GAPDH panel correspond mutant GAPDH-K225A which is HA-tagged. The figure presents reformatted images from the original immunoblots in **Fig. 6d** for easier interpretation. Lanes were rearranged to match the order shown in **Fig. 6e**.

**Supplementary Table 1: Clinical characteristics of human subjects for myocardial immunofluorescence analysis.**

Myocardial tissues were obtained from control patients who had undergone heart transplantation (HTx) and patients with dilated hypertrophic cardiomyopathy (dHCM). BNP, B-type natriuretic peptide; LVEF, left ventricular ejection fraction.

| **Control** | **Case #1** | **Case #2** | **Case #3** |
|---|---|---|---|
| Age (years) | 43 | 43 | 61 |
| Sex | Male | Male | Male |
| LVEF (%) | 60 | 66 | 61 |
| BNP (pg/mL) | 51.1 | 64.5 | 92.4 |
| Pre-HTx diagnosis | Endocardial fibroelastosis | Anthracycline-induced cardiomyopathy | Dilated cardiomyopathy |
| Donor age | 30s | 50s | 30s |
| Donor sex | Female | Male | Male |
| Cause of brain death | Traumatic injury | Post-cardiac arrest brain injury | Subarachnoid haemorrhage |

| **dHCM** | **Case #4** | **Case #5** | **Case #6** |
|---|---|---|---|
| Age (years) | 46 | 49 | 53 |
| Sex | Male | Female | Male |
| LVEF (%) | 15 | 22 | 30 |
| BNP (pg/mL) | 172.2 | 1336 | 357.5 |

**Supplementary Table 2: In vitro screening for off-target activities of RR.**

RR exhibited no significant interaction with primary targets, including receptors, channels, transporters, and enzymes, at a concentration of 1 μM, which is 1000-fold higher than the concentration required to inhibit GAPDH-Siah1 binding.

| Primary target | % inhibition |
|---|---|
| **Adenosine $A_1$** | 1 |
| **Adenosine $A_{2A}$** | 8 |
| **Adrenergic $\alpha_{1A}$** | 9 |
| **Adrenergic $\alpha_{1B}$** | -3 |
| **Adrenergic $\alpha_{2A}$** | 12 |
| **Adrenergic $\beta_1$** | 0 |
| **Adrenergic β2** | 2 |
| **Calcium Channel L-Type, Dihydropyridine** | -4 |
| **Cannabinoid $CB_1$** | 9 |
| **Dopamine D1** | 4 |
| **Dopamine D2S** | 4 |
| **GABAA, Flunitrazepam, Central** | -3 |
| **GABAA, Muscimol, Central** | 1 |
| **Glutamate, NMDA, Phencyclidine** | 1 |
| **Histamine $H_1$** | 0 |
| **Imidazoline $I_2$, Central** | 9 |
| **Muscarinic $M_2$** | -2 |
| **Muscarinic $M_3$** | 5 |
| **Nicotinic Acetylcholine** | -3 |
| **Nicotinic Acetylcholine α1, Bungarotoxin** | 3 |
| **Opiate μ (OP3, MOP)** | 4 |
| **Phorbol Ester** | 7 |
| **Potassium Channel [$K_{ATP}$]** | -10 |
| **Potassium Channel hERG** | -2 |
| **Prostanoid $EP_4$** | 3 |
| **Rolipram** | 7 |
| **Serotonin (5-Hydroxytryptamine) 5-$HT_{2B}$** | 19 |
| **Sigma $\sigma_1$** | 7 |
| **Sodium Channel, Site 2** | 14 |
| **Transporter, Norepinephrine (NET)** | 17 |

**Supplementary Table 3: Nuclear GAPDH-associated partner proteins involved in chromatin remodelling.**

Among the 567 potential interactors of nuclear GAPDH identified by LC-MS/MS, GO analysis ranked "chromatin remodeling" (GO:0006338) as the top term based on the number of associated proteins. The 56 proteins classified under this term are listed.

| | | | | | | |
|---|---|---|---|---|---|---|
| ACP1 | DDX17 | HDAC1 | MCM6 | PPM1A | RPS6KA3 | SMARCC2 |
| ACTB | DDX39B | HDAC2 | MCM7 | PPP1CA | RUVBL1 | SMARCD1 |
| ACTL6A | DHX9 | HUWE1 | MTOR | PPP2CB | RUVBL2 | SMG1 |
| ARID1A | DHX15 | ILKAP | MTREX | PRMT1 | SETD3 | SNRNP200 |
| CARM1 | EIF4A1 | KDM5C | NAA50 | PRMT5 | SF3B1 | UCHL5 |
| CDK1 | EIF4A3 | MAP2K1 | NEK9 | RBBP4 | SFPQ | VCP |
| DDX1 | GTF3C4 | MCM3 | NSF | RBBP7 | SKP1 | VPS4B |
| DDX6 | HCFC1 | MCM4 | PKM | RECQL | SMARCA4 | WNK1 |
